# Cross-chemistry single-nucleus RNA-seq identifies gene length and CpG-island promoters as determinants of transcriptional noise

**DOI:** 10.64898/2026.08.25.747056

**Authors:** Rafal Czapiewski, Michael Chiang, James Ding, Catherine Naughton, Graeme R. Grimes, Davide Marenduzzo, Nick Gilbert

## Abstract

Cell-to-cell transcriptional heterogeneity, or noise, is an intrinsic property of the transcriptome with implications for development, disease progression, and aging. Bulk RNA-seq masks this variability by averaging gene expression across cells, whereas single-cell RNA sequencing (scRNA-seq) resolves it. Nevertheless, separating biological noise from technical variance remains challenging, particularly across platforms with different chemistries. We benchmarked two widely adopted technologies, Evercode WT (SPLiT-seq, Parse Biosciences) and Chromium (10x Genomics), on human lymphoblastoid nuclei. Evercode WT achieved targeted sequencing depth and nuclei number far more reliably, and its random-hexamer priming yielded more intronic reads and non-coding RNA genes; Chromium recovered more cells and detected polyadenylated transcripts and cell-line markers more sensitively. Despite these opposing biases, the platforms showed comparable gene detection and strongly correlated expression profiles. Using datasets from both platforms, we defined a noise metric detrended from mean expression and showed that per-gene estimates were reproducible across chemistries. Noise was lower in G2M than in G1 and was most strongly associated with gene length rather than exonic length. Expression of genes with CpG-island promoters was less variable than that of those without. This study establishes a platform-independent basis for quantifying transcriptional noise and a framework for selecting an appropriate scRNA-seq platform.

**Highlights:**

- Parse Evercode WT demonstrates superior predictability in targeted cell recovery and sequencing depth estimations compared to 10x Chromium.
- Distinct biases: Parse captures intronic sequences; 10x targets polyadenylated mRNA.
- Detrended transcriptional noise is reproducible across both barcoding chemistries.
- Noise scales with gene length, not exonic length, and is lower at CGI promoters.

## Introduction

Single-cell (sc) and single-nucleus (sn) RNA sequencing (scRNA-seq, snRNA-seq) are currently standard techniques in molecular biology research for assessing gene expression. Unlike bulk RNA-seq, scRNA-seq detects variations in gene expression between individual cells, enabling identification of cell types, single-cell states, and transcriptional heterogeneity within cell populations. Since the first single-cell transcriptome experiment reported by Tang *et al.* in 2009 (1), various adaptations of the technology have been applied to numerous cell lines and to primary cells obtained from dissociated tissues across different species (2). The primary achievement of single-cell platforms to date is arguably the creation of tissue cell atlases - indispensable resources in biomedicine- alongside the discovery of several new cell types, such as stem cells in oligodendroglioma (3), respiratory airway secretory cells (4), thymus M cells (5), and EndoMac cells, which are aortic endothelial progenitors (6). Single-cell approaches are also crucial for translational research, as diseases often present with abnormal cell type proportions within tissues, such as a reduction in a subclass of neurons in Alzheimer’s Disease (7), depletion of a subtype of intraepithelial lymphocytes in celiac disease (8), and an increase in type I fiber myonuclei in muscular dystrophy (9).

Furthermore, single-cell techniques are applicable to studies of transcriptional states during development and aging, as well as transcriptional heterogeneity (10–12). For example, they enabled the discovery of new transcriptional signatures of T cells originating from different tissues (13) and the identification of heterogeneity among these macrophages in the tumor environment (14,15). In addition, the development of single-cell transcriptome platforms has enabled other techniques such as scATAC-seq (16), scHi-C (17,18), scChIP-seq (19), and scDamID (20,21), complemented by the emergence of *spatial transcriptomics* based on *in situ* hybridization (22). Many techniques are now combined in *multiome* setups, enabling, for example, gene expression analysis and chromatin accessibility in a single cell (23). The growing number of scRNA-seq methods necessitates systematic comparisons and benchmarking for data validation. Furthermore, identifying platform-specific bias is crucial, as each chemistry and transcript barcoding method may introduce bias based on gene length, GC content, or RNA localization (24,25).

In principle, all single-cell transcriptomic techniques rely on fusing a unique barcode (a short synthetic oligonucleotide DNA sequence) to mRNA molecules from individual cells. Microfluidic systems can deliver these barcodes by enclosing a single cell within a bead, oil droplet, or gel. Microcapsules carry unique barcodes, enabling the assignment of each transcript to a unique cell after sequencing. In non-microfluidic platforms, barcodes are delivered by placing cells in liquid solutions in individual microtubes or multi-well plates, where cells undergo permeabilization or other forms of lysis to deliver the barcodes (26,27). Recent studies systematically compared major technologies using different barcoding workflows across several cell lines and tissues at the single-cell and single-nucleus level, aiming to identify the platform that processes the most cells per sample while detecting the highest number of unique transcripts and expressed genes (28–30). At the same time, the goal is to avoid duplicates (also called doublets), i.e., two or more cells with the same sets of barcodes. Importantly, bioinformatics can remove doublets somewhat efficiently from datasets (31). Because of its robust performance, the most popular scRNA-seq platform is currently a microfluidic platform, Chromium from *10x Genomics*. This microfluidic method uses gel beads with barcodes, and each bead captures one cell. During reverse transcription, barcodes are fused to mRNA, creating a unique *single-cell* signature and allowing assignment of each barcoded cDNA library to an individual cell (**Supplementary Fig. S1A**). However, 10x Chromium comes with its own limitations. Although it produces high-quality data, 10x Chromium is expensive for many labs, requires costly equipment, and, perhaps most importantly, detects a low number of genes. At the other end of the price spectrum in single-cell technologies, Parse Bioscience commercialized the SPLiT-seq method and developed a single-cell RNA-seq kit branded Evercode WT based on combinatorial transcript barcoding (27). This platform takes advantage of split-pool ligation-enabled combinatorial barcoding (**Supplementary Fig. S1A**).

In this study, we assess the 10x Chromium and Evercode WT single-nucleus RNA sequencing platforms by using GM12878 lymphoblastoid nuclei. Additionally, we carried out a whole-cell single-cell RNA sequencing (scRNA-seq) experiment on the same cells using the 10x Chromium system. We compare the efficiency of the two platforms, examine batch effects, assess sequencing depth, and investigate reproducibility across both platforms. We report a biological detection bias caused by differences in reverse-transcription chemistry between the two platforms. We benchmark transcriptional heterogeneity across technologies and provide a platform-independent measure of transcriptional noise. We also measure noise throughout the cell cycle and find that G2M is the phase that shows the least transcriptional noise. Finally, we use the datasets obtained to examine the gene characteristics that might drive transcriptional noise. Of the gene features that we tested, gene length is the strongest positive predictor of transcriptional noise. Importantly, we observed this heterogeneity independently of the technology used and regardless of the chemistries applied during reverse transcription, the first step in barcoding the transcripts. We also note that transcription start sites within a CpG island are less noisy than those outside it. Using these noise parameters, we systematically analyze how transcriptional heterogeneity varies with cell-cycle phase, gene length, and promoter architecture, establishing a robust quantitative basis for understanding the biological factors that drive expression noise.

## Results

### Experimental design and comparison of sequencing libraries

To evaluate the applicability of Parse Evercode WT and 10x Chromium scRNA-seq technologies for studying transcriptional heterogeneity, we aimed to compare how each platform contributes to cell-to-cell variation in gene expression and to test the reproducibility of results from technical replicates. To measure efficiency metrics for both platforms, we used isolated nuclei from GM12878 cells. All experiments involved four technical replicates to account for variability in sample preparation and processing. Additionally, we included a relatively higher number of replicates to minimize technical noise, thus improving the precision and reliability of the results. Besides single-nucleus experiments, we conducted one whole-cell experiment using scRNA-seq on a 3’ 10x Chromium platform as a reference (**Fig. 1A**). The main difference between Parse Evercode WT and 10x Chromium lies in their transcript barcoding methods and resulting library structures (**Fig. 1B** and **Supplementary Fig. S1A** and **S1B**). Both platforms’ libraries include standard Illumina indexing (i5 and i7) and unique platform-specific cell and sample identification barcodes (Parse WT Evercode: 48 bp vs. 10x Chromium: 28 bp). Although the barcode lengths differ, which suggests more “usable reads” from the 10x platform, the UMI and Poly(dT) sequence lengths balance this difference, leading to similar sequencing yields (**Fig. 1C**).

**Figure 1.**
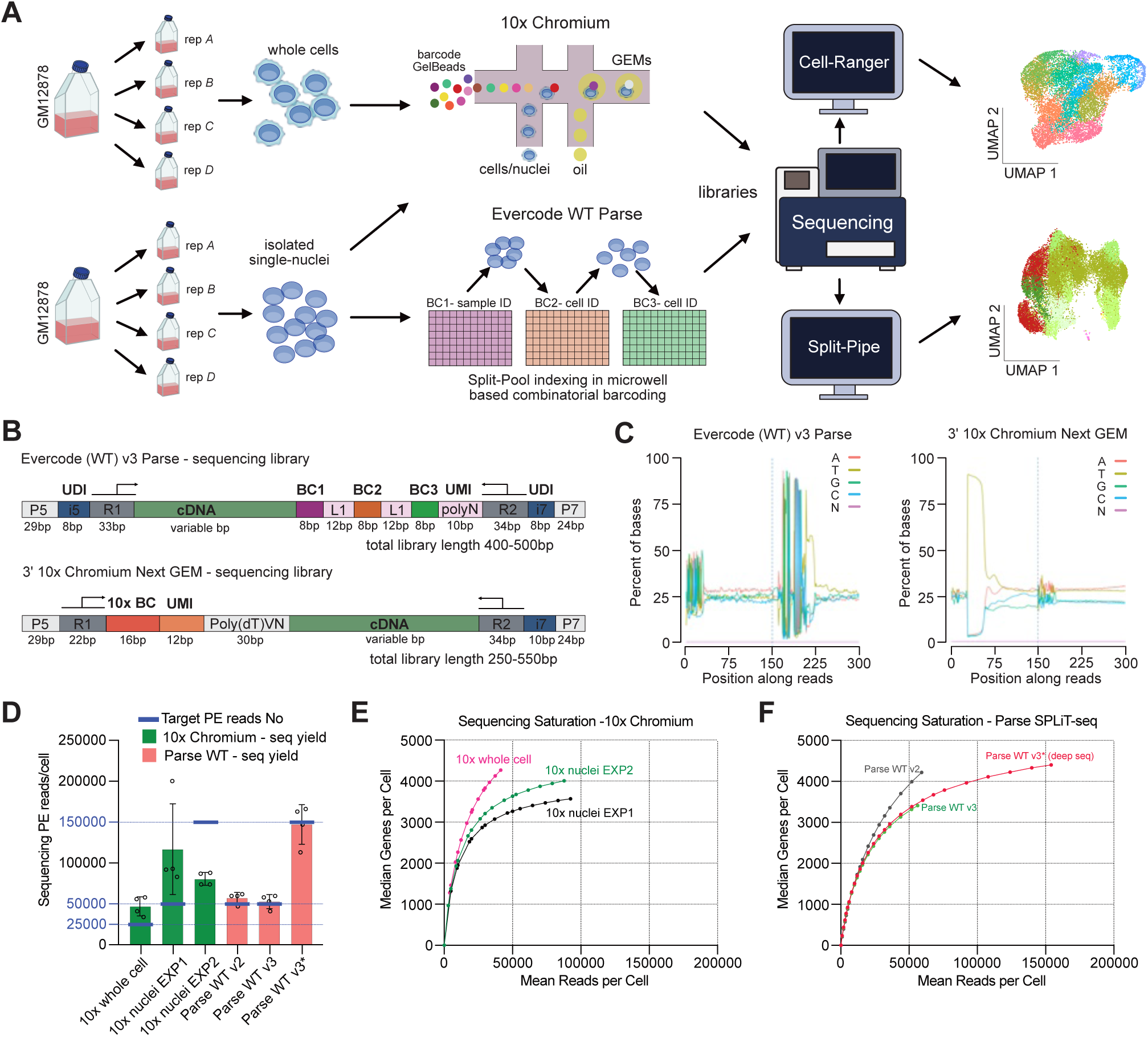
Experimental workflow and primary QC comparison. **(A)** Each experiment includes four technical replicates (separate cell culture flasks). In 10x Chromium microfluidic technology, single cells are captured with a bead saturated with barcodes inside an oil droplet. Evercode WT Parse uses sequential combinatorial barcoding in a split-pool process. Libraries are sequenced, and results are analyzed with Cell-Ranger (10x) and Split-Pipe (Parse). See also Figure S1 A. **(B)** Schematic of sequencing libraries. BC1 - sample identification barcode; BC2-BC3 - single-cell identification barcodes. UMI - unique molecular identifier. 10x BC - Chromium 10x barcode for sample and single-cell ID. R1 and R2 are sequencing read primers. Primers i5 and i7 - Illumina demultiplexing index primers. **(C)** Representative plots of sequencing data. **(D)** Prediction of sequencing depth. Average sequencing yield (paired-end reads) per cell in each experiment (green bars – 10x, red bars – Parse), plotted against targeted sequencing depth - blue dashed bars. Circles denote technical replicates, and error bars show SD. **(E** and **F)** Sequencing saturation. Median number of genes per cell at down-sampled sequencing depth. Representative curves from one sequencing library per experiment are shown for each platform. All technical replicates are shown in **Supplementary Fig. S1H** and **S1I**. An asterisk (*) indicates a Parse WT v3 experiment with increased read depth (50K reads/cell vs. 150K reads/cell). See also **Supplementary Table S1.**

### Sequencing depth

Choosing sequencing depth is the main challenge when planning single-cell RNA-seq experiments. Both platform manufacturers recommend at least 20,000 paired-end (PE) reads per cell, but there is no guidance for isolated nuclei. Assuming fewer transcripts are available in nuclei, we targeted depths of 25,000 reads per cell for the 10x whole-cell experiment, and two depths for nuclei: 50,000 and 150,000 reads per nucleus (**Fig. 1D** and **Table 1**). Surprisingly, the predictability of sequencing yield was a major difference between platforms. Parse Evercode WT experiments achieved nearly 100% targeted sequencing depth. This high predictability of sequencing yield from the Parse platform was also consistent across all replicates and experiments with different targeting depths, i.e., 50,000 or 150,000 PE reads per cell (**Fig. 1D**).

**Table 1.** Summary of the experimental yield. Six single-cell RNA-seq experiments were conducted, each with four technical replicates: **A, B, C,** and **D. Target nuclei No./sample –** refers to the estimated number of cells and nuclei detected per technical replicate. Six experiments were performed, each with four independent technical replicates (see also **Figure 1** for the experimental setup). **Target seq reads** – indicate the expected sequencing depth (see also **Table S1** for a detailed comparison of targeted and yield sequencing depth across experiments and platforms). **\*\*<u>sc</u>**RNA-seq 1 Ox Chromium is a “whole cell” experiment, so the number of cells is provided. **Genes/cell** – represents the number of individual genes detected in a replicate per cell. **Nuclei No./sample** - number of cells detected in each replicate and experiment. Total nuclei - all cells detected per experiment. Refer also to **Table S1** for technical details on sequencing and sequence read yield.

| Platform/Exp | scRNA-seq**<br>10x Chromium | snRNA-seq<br>10x Chromium<br>EXP1 3' v3.1 | snRNA-seq<br>10x Chromium<br>EXP2 3' v3.1 | snRNA-seq<br>Evercode (WT)<br>Parse v.2 | snRNA-seq<br>Evercode (WT)<br>Parse v.3* | snRNA-seq<br>Evercode (WT)<br>Parse v3* deep |
| --- | --- | --- | --- | --- | --- | --- |
| Targeted nuclei<br>No./sample | 10,000** | 8,000 | 10,000 | 5,000 | 5,000 | 5,000 |
| Targeted<br>reads/nuclei | 50k/cell | 50k/nuclei | 150k/nuclei | 50k/nuclei | 50k/nuclei | 150k/nuclei |
| Genes/cell | A - 614 | A - 2900 | A - 3851 | A - 4264 | A - 3550 | A - 4616 |
|  | B - 4924 | B - 3698 | B - 4006 | B - 4198 | B - 4092 | B - 5125 |
|  | C - 4272 | C - 3571 | C - 3781 | C - 3692 | C - 2380 | C - 3276 |
|  | D - 3848 | D - 3100 | D - 3818 | D - 4295 | D - 3565 | D - 4624 |
| Nuclei No./<br>sample | A - 4315 | A - 1188 | A - 11831 | A - 4352 | A - 5887 | A - 5863 |
|  | B - 6361 | B - 3398 | B - 9993 | B - 4398 | B - 5259 | B - 5232 |
|  | C - 8590 | C - 3428 | C - 11591 | C - 5500 | C - 6145 | C - 6135 |
|  | D - 8941 | D - 4069 | D - 10966 | D - 3851 | D - 5632 | D - 5602 |
| Total nuclei | 28,207 | 12,083 | 44,381 | 18,013 | 22,923 | 22,832 |
| Total PE reads | 1,366.8 M | 1,363.2 M | 3,602.3 M | 1,036.0 M | **1,222.4 M** | **3,352.7 M** |

In contrast, the 10x Chromium platform consistently failed to reach the target sequencing depths. The 10x whole cell experiment targeted 25,000 reads per cell, and the 10x nuclei EXP1 targeted 50,000 reads per nucleus; both produced roughly double the intended depth of around 50,000 reads per cell and 100,000 reads per nucleus, respectively. In the 10x nuclei EXP2, however, we achieved only half of the targeted depth, possibly due to an unintended overload of nuclei per sample (**Fig. 1D**). Therefore, technical differences between the two platforms cause discrepancies in sequencing predictability. In Parse, however, after three rounds of split-pool barcoding, nuclei (or cells) are counted just before lysis, the first step of sequencing library preparation, ensuring an exact number of cells are lysed into the library prep. In contrast, 10x Chromium platforms allow cell counting only before loading cells into a microfluidic device, where they are barcoded and lysed, preventing accurate prediction of the number of cells per library. Ultimately, precise sequencing depth prediction requires knowing the exact number of cells per library (**Supplementary Fig. S1A;** see also the Methods section).

Predicting sequencing depth is essential for maximizing data output and estimating sequencing saturation, the point in a DNA sequencing experiment when increasing the number of reads no longer yields more unique UMIs and genes. As expected, the 10x whole-cell experiment yielded the highest number of detected genes because transcripts are more abundant in the cytoplasm. However, the saturation curve indicates that doubling or even tripling the read depth could help detect additional new genes from whole cells (**Fig. 1E**). Sequencing of 10x nuclei EXP1 and EXP2 is closer to saturation, and we estimate that 100,000 reads per nucleus would be optimal for these experiments; however, predicting the exact depth for 10x experiments remains challenging. Importantly, we saw similar saturation levels in Parse, showing that 50,000 reads per nucleus nearly double the number of genes detected compared to the recommended 20,000 reads. Furthermore, in both platforms, at 50,000 reads, approximately 3,500 genes are detected per nucleus, indicating similar library efficiency for both technologies. Finally, saturation curves suggest that sequencing between 100,000 and 150,000 reads per nucleus maximizes yield on both platforms (**Fig. 1E** and **1F**, see also **Supplementary Fig. S1H** and **S1I** for replicate data).

### Platform-specific bias

Beyond the predictability of sequencing yield, both platforms differ in their molecular biology approach used for utilizing mRNA templates in cDNA synthesis. Specifically, 3’ 10x Chromium uses oligo(dT) primers for barcode fusion via reverse transcription. In contrast, Parse SPLiT-seq employs both random hexamers and oligo(dT) primers in the first barcoding reaction (**Supplementary Fig. S1A** and **S1B**; see also the Material and Methods section). In theory, this difference in cDNA synthesis could introduce bias, since 3’ oligo(dT) primers would anneal to polyadenylated transcripts, while random hexamers would detect any RNA transcribed in the cells. As predicted, we observe a 3’ bias in gene body coverage from the 10x Chromium platform. Interestingly, although we expected uniform gene coverage from Parse, we observe some 5’ bias, possibly due to the detection of short, incomplete transcripts (**Fig. 2A**). This bias affects the distribution of reads along gene bodies. 10x Chromium produces more reads from 3’UTR exons than Parse, which in turn shows more intronic reads in our experiments (**Fig. 2B**). A similar pattern appears in stacked read coverage, showing uniform distribution of reads mapping to all exons in Parse, whereas most reads in the 10x experiment map mainly to the 3’ exon. Examples are shown from genes *HLA-B* and *MYC* (**Fig. 2C**).

**Figure 2.**
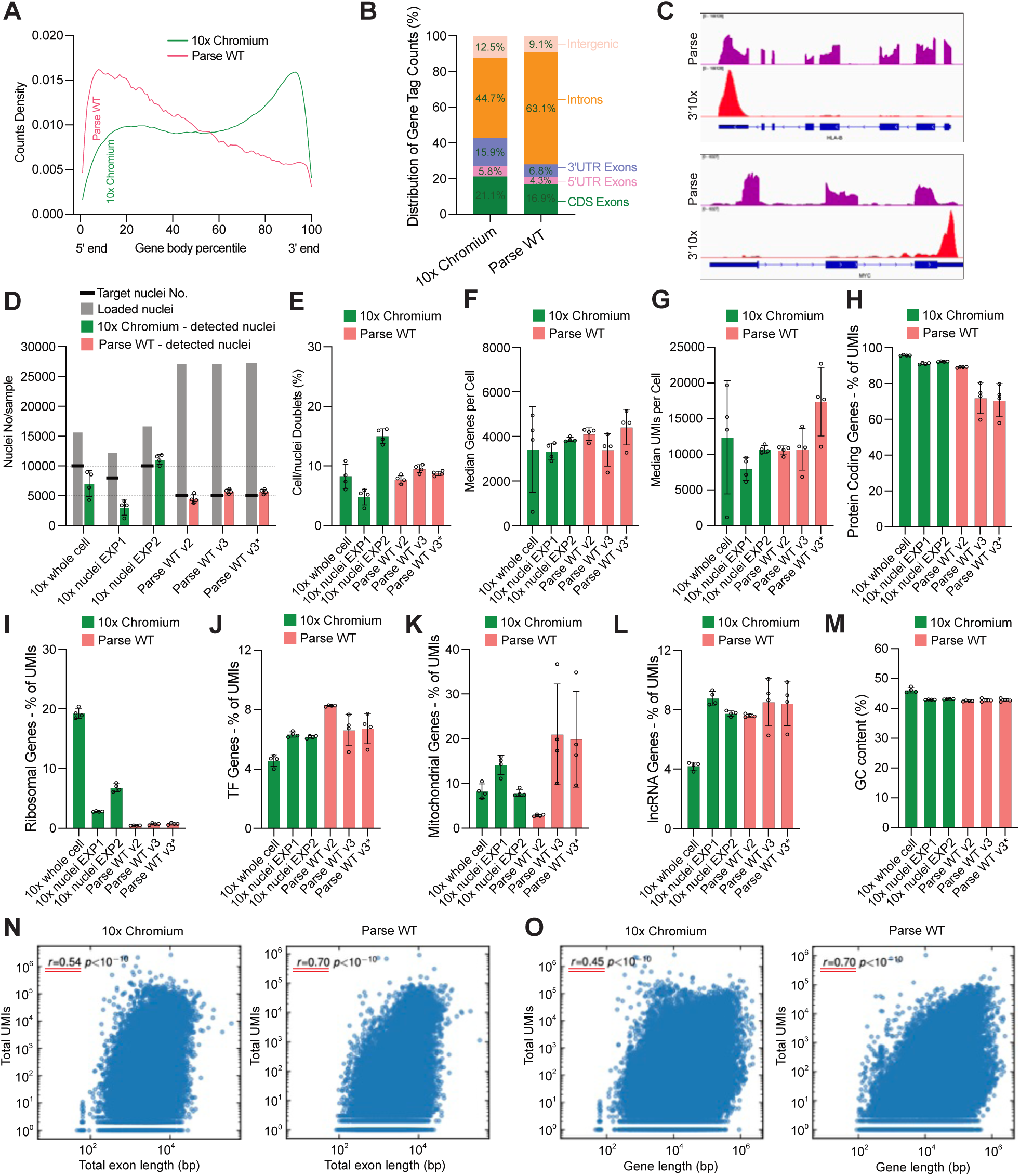
Sequencing library efficiencies. **(A)** Gene coverage is shown as a plot of counts density against normalized gene body length from 5’ to 3’. Parse WT includes four technical replicates from the experiment “Parse WT v2,” and the 10x experiment has four replicates from “10x nuclei EXP2.” **(B)** Distribution of reads across gene body features—including CDS exons, 5’ UTR exons, 3’ UTR exons, introns, and intergenic regions—shown as a proportion of gene tag counts. **(C)** Stacked snRNA-seq traces for both platforms display read coverage of HLA-B and MYC genes, with 3’ enriched reads for the 10x platform and exon-enriched reads for Parse WT. **(D)** Nuclei retention comparison. Grey bars show the number of loaded cells per replicate; green and red bars represent the average detected cell numbers, and circles indicate technical replicates. Black dashed bars show the targeted number of nuclei. **(E)** Cell doublet rates detected by *scDBL_finder* software; circles represent individual replicate values, bars show mean values, and error bars indicate SD. (**F** and **G)** Bar graphs display median gene and UMI counts per cell, respectively. Circles show replicate data; error bars represent SD. **H-L.** Gene detection by platform. Each graph shows the percentage of UMIs mapped to protein-coding genes (H), ribosomal protein genes (I), transcription factor genes (J), mitochondrial genes (K), and lncRNA genes (L). **(M)** Percentage of GC nucleotides in detected genes. **(N** and **O)** Platform-specific bias in exon length (N) and gene length (O) is shown as scatterplots of gene length versus the number of UMIs, where *r* indicates the correlation between gene length and expression for all genes. See also Table S2 and Figure S2.

Another parameter obtained from sequencing data is the number of cells per sample. Previous reports suggest that 10x Chromium has better cell retention than Parse (30). We observed a similar trend with isolated nuclei. 10x Chromium achieved 65% recovery in 10x nuclei EXP2; however, other 10x experiments show 30-40% retention. Using the same counting method, we found that Parse, as previously reported, retains about 20% of cells (**Fig. 2D** and **Supplementary Fig. S2B**). However, it is important to note that these calculations might not present the full picture. We observe that during the barcoding step in Parse, indeed approximately 20-50% of nuclei are lost (**Supplementary Fig. S2C**). This measurement is possible because, in Parse, barcoding occurs in permeabilized but intact nuclei, which enables counting. However, we account for inflated cell loss here because, in the next step of the protocol, only a fraction of the nuclei obtained from barcoding is used for cell lysis for reaction in library preparation (**Supplementary Fig. S1A**). In our Parse experiment, we used 20,000 cells. With this additional counting step, we see little to no loss of nuclei after sequencing, detecting 18,000 with Parse v.2 and 22,000 nuclei with Parse v.3 (**Table 1** and **Supplementary Fig. S2C**). These different ways of counting cells complicate direct comparisons between platforms; nevertheless, they suggest that Parse indeed requires more starting material for barcoding than 10x Chromium. However, the cell-counting step just before lysis allows precise cell-number loading for Parse libraries. This enables strong predictability of the cell number detected and achieves nearly 100% of the targeted cell number in Parse (**Fig. 2D** and **Table 1**). This results in nearly 100% predictability of sequencing yield depth for Parse (**Supplementary Fig. S1H**).

For further quality control, we calculated the cell doublet ratio (i.e., cells sharing the same set of barcodes) using the bioinformatics tool *scDblFinder* (32). Our results indicated that Parse experiments had a similar doublet rate compared to 10x Chromium, with approximately 8-10% of cells classified as doublets. However, in 10x nuclei EXP2, we identified roughly 15% doublets, possibly due to library overloading (**Fig. 2E**). This, however, cannot be precisely predicted in 10x, and it’s a general reminder that overloading the system increases the doublet rates.

Current reports on the sensitivity of single-cell platforms suggest that Parse has a slight advantage in detecting unique transcripts compared to the 10x Chromium. However, these findings vary across the literature (30,33,34). We aimed to compare these sensitivity metrics in isolated nuclei from GM12878 cells. We found that both platforms detect a similar number of genes per nucleus and show comparable variation between replicates (**Fig. 2F** and **Table 1**). Similarly, the number of UMIs is consistent across experiments, except for Parse WT v3*, which involves sequencing libraries at greater depth. Surprisingly, the 10x whole-cell experiment did not yield significantly more unique genes or UMIs than the single-nuclei experiments, but it displayed large variation between technical replicates (**Fig. 2F** and **2G**).

To further dissect platform differences and potential bias, we analyzed the proportions of detected gene types. Interestingly, 10x Chromium detected a slightly higher ratio of protein-coding genes than the Parse experiments. Both the 10x whole-cell and 10x single-nucleus experiments showed that over 90% of genes are protein-coding. However, many genes from the whole-cell experiments encoded ribosomal proteins, while this fraction was lower in the 10x single-nucleus samples (**Fig. 2H**). Importantly, as previously observed (30,34), Parse detected a very small proportion of ribosomal genes, a phenomenon not fully understood but previously reported (30) (**Fig. 2I**, **3B** and **3D**).

We also analyzed transcription factor gene levels and found no significant differences between platforms in the single-nucleus samples (**Fig. 2J**). Next, as a biological quality check, we examined the percentage of mitochondrial genes among all UMIs detected and observed similar values of around 8-12% for 10x experiments. However, a clear difference emerged between v2 and v3 Parse experiments. With less than 5% mitochondrial genes in v2 and about 20% in Parse v3, we suspect that lower-quality nuclei were loaded for barcoding in the latter. The large variation between samples in v3 highlights the need for additional quality control and underscores the importance of technical replicates (**Fig. 2K** and **Supplementary Fig. S2D**).

Long non-coding RNAs were also proposed as another indicator of sample quality in scRNA-seq experiments (35). In our study, both platforms showed a very similar proportion of lncRNA among UMIs at about 8% in single-nucleus experiments. Interestingly, the whole-cell 10x experiment showed only 4%, close to previous reports (33) (**Fig. 2L**).

All the parameters mentioned above could be explained by genetic factors such as GC content or gene length, as suggested in previous studies (30,34). Interestingly, we did not observe differences in GC content between Parse and 10x, as reported in studies conducted on whole cells (30). In our experiments, both platforms showed a GC content of approximately 42% in isolated nuclei, whereas the 10x whole-cell experiment showed 45% (**Fig. 2M**). This is close to the reported human genome average of 41% (36). The key difference between the two technologies, however, is gene-length bias. As described earlier, differences between random hexamers and oligo(dT) contribute to gene body coverage bias (**Fig. 1A** and **1C**). Therefore, we examined whether differences in reverse transcription chemistry introduce length-detection bias across exons and genes. Consistent with previous studies, Parse Evercode WT showed a pronounced preference for longer exons (**Fig. 2N**) and longer genes (**Fig. 2O**), indicating that combining random hexamers and oligo(dT) primers drives gene-length bias in the Parse platform.

### Biological parameter yields vary between Parse Evercode WT and 10x Chromium

QC parameters from experiments on both platforms confirm that 10x Chromium is biased toward polyadenylated transcripts. At the same time, Parse Evercode WT shows a preference towards longer genes due to the difference in reverse transcription chemistry. These differences in chemistry and resulting biases suggest that both platforms may yield different biological metrics in single-nucleus transcriptome studies. To examine this, we first compared the expression of selected genes across various categories, including housekeeping, ribosomal, and lncRNA genes. We found that 10x nuclei EXP1 and EXP2 showed greater sensitivity for *GAPDH* and *ACTB* genes than Parse Evercode WT. However, both platforms were similarly effective at detecting *CD47* and *B2M* gene expression (**Fig. 3A**). Interestingly, both platforms detected the chromatin-organizing genes *CTCF* and *YY1* at similar levels (**Fig. 3B**). The genes most differentially expressed between platforms were lncRNA *TALAM1,* which was highly expressed in Parse but nearly undetectable in 10x nuclei, and lncRNA *SNHG3*, which was highly expressed in 10x experiments but minimally detected by Parse (**Fig. 3B, Supplementary Fig. S3A** and **S3B**). This difference likely reflects that *TALAM1* is not polyadenylated, whereas *SNHG3* produces polyadenylated transcripts, making platform-bias interpretation for long non-coding RNAs more complex. As explained earlier, oligo(dT) used in the 10x platform and random hexamers in Parse introduce bias linked to polyadenylation status (**Fig. 3C**). Intriguingly, as previously shown, ribosomal genes were more expressed in 10x experiments than in Parse (**Fig. 3B**). Perhaps it could be expected that Parse, which uses oligo(dT) and random hexamers, would be more sensitive to these multi-exonic and polyadenylated transcripts; however, this is not the case in our study and in published reports (**Fig. 3D**). One possible explanation is that ribosomal protein-coding genes, which have a higher GC content (∼49%) than the genome-wide average (∼41%), would be more detectable in 10x if bias toward higher GC content genes existed. Nonetheless, we did not observe the previously reported GC content bias in the technologies used (**Fig. 2M**). Therefore, the cause of the discrepancy in ribosomal gene detection remains unclear (30,34). Next, we examined the expression of cell-type markers. GM12878 is an adult human lymphoblastoid cell line (LCL) derived from a female of European ancestry. LCLs are generated by transforming peripheral blood lymphocytes with Epstein-Barr Virus (EBV), which immortalizes resting B cells and yields an immortal, actively proliferating B cell population (37). Human B cells are diverse, with multiple subsets, and express core markers such as CD19, IgM, IgD (*IGHD*), CD27, and CD38. LCLs are also negative for the NK cell marker CD56 (coded by *NCAM1*) and the T cell marker CD3 (coded by *CD3E*, *CD3D,* and *CD3G*) (38). However, to our knowledge, no study has fully validated these markers for B cells and immortalized LCLs (39). In our experiments, both platforms accurately captured both positive and negative markers expression profiles as expected. Nonetheless, 10x showed high expression of classic positive LCL markers such as *IGHD*, *CD79A*, and *CD81*, while Parse detected these markers in fewer cells and at lower levels. This potentially increases 10x relevance to cell biology by more accurately reflecting underlying biological states (**Fig. 3E**). Importantly, both platforms detected minimal levels of negative markers: *CD3E*, *CD3D,* or *NCAM1* (**Fig. 3E**).

**Figure 3.**
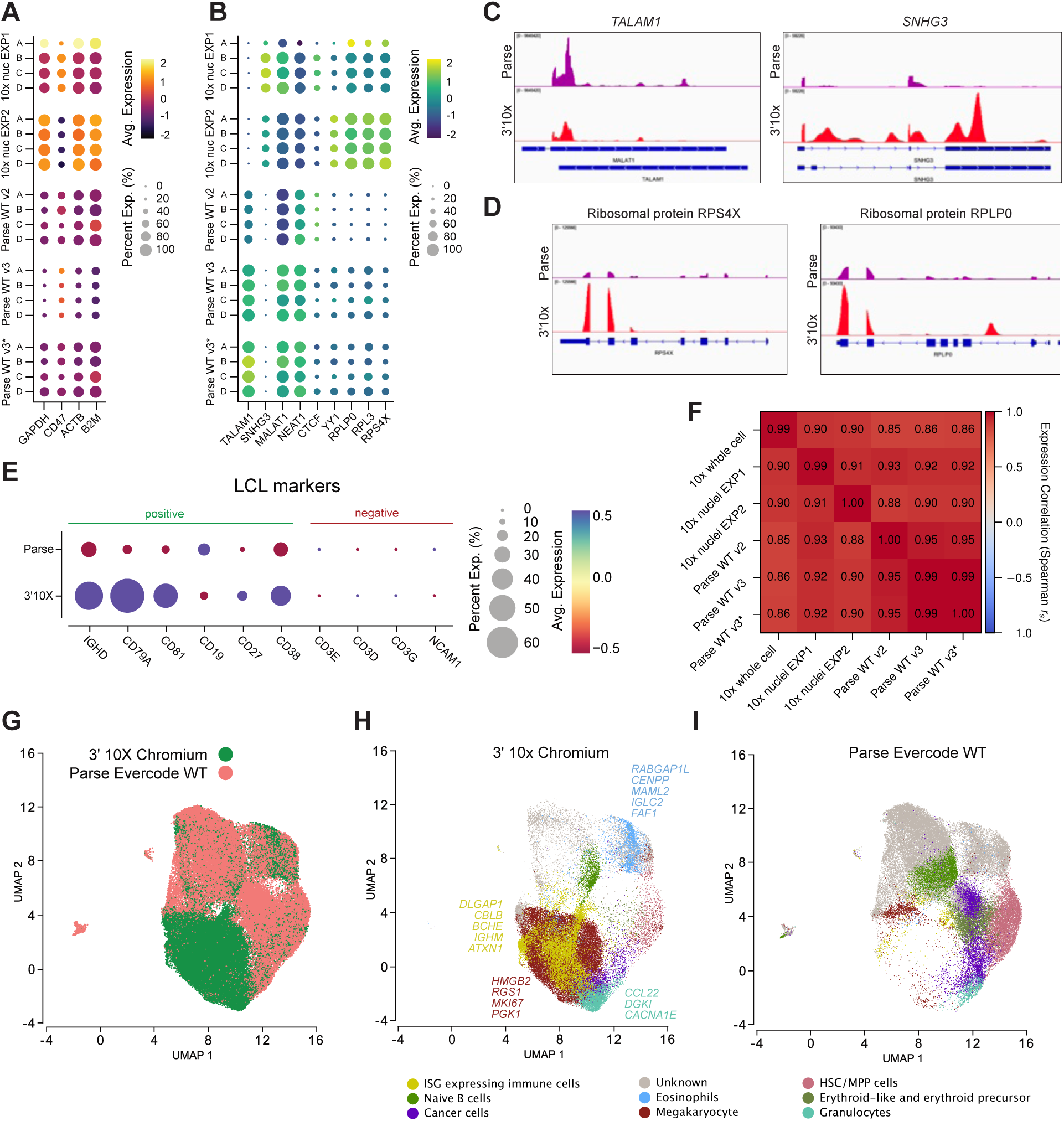
Platform-dependent variations in biological metrics. **(A** and **B)** Dot plots depicting the expression of (A) housekeeping genes and (B) selected functional gene categories, including lncRNAs (*TALAM1*, *SNHG3*, *MALAT1*, *NEAT1*), chromatin regulators (*CTCF*, *YY1*), and ribosomal proteins (*RPLP0*, *RPL3*, *RPS4X*) across technical replicates (A, B, C, and D) for both platforms. Circle size represents the percentage of expressing cells (Percent Exp.), and color intensity indicates mean expression levels. **(C** and **D)** Representative pseudobulk snRNA-seq genomic coverage tracks comparing 10x Chromium and Parse Evercode WT for (C) lncRNAs (TALAM1, SNHG3) and (D) ribosomal protein-coding genes (RPS4X, RPLP0). **(E)** Dot plot displaying the expression of canonical positive and negative marker genes in lymphoblastoid cell lines (LCLs). **(F)** Pairwise Spearman correlation matrix (*r_s_*) of mean gene expression across experimental batches and platforms. **(G)** UMAP projection of single nuclei colored by platform (10x Chromium, green; Parse Evercode WT, coral). **(H** and **I)** UMAP visualization showing Trailmaker-annotated cell types and clusters separately for (H) 10x Chromium and (I) Parse Evercode WT. Annotated cell-type clusters and representative marker genes are color-coded according to the legend.

To further compare the two platforms biologically, we examined the similarity of their bulk transcriptomic profiles. We observed a strong correlation between platforms and replicates in gene expression levels (**Fig. 3F**). Notably, Parse Evercode WT shows slightly higher correlation and reproducibility across experiments, with Spearman’s *r*_s_ >0.95. Furthermore, for technical replicates, Parse also shows superior reproducibility compared with 10x. In contrast, 10x showed poorer consistency between replicates, and the 10x whole-cell experiment showed the highest variation between technical replicates (**Supplementary Fig. S3B**).

Because the heterogeneity of GM12878 cells remains poorly characterized, we evaluated signature gene sets associated with known cell types. To test clustering of distinct subpopulations, we applied batch correction by *Harmony* and dimensionality reduction using the *Trailmaker* platform (40). We found that cells mostly cluster by platform, with limited overlap, as shown in the UMAP projection plots (**Fig. 3G**). This suggests a significant platform-specific bias, likely driven mainly by the reverse transcription chemistry and physical cell handling on both platforms.

To further evaluate detection of biological markers in cell subpopulations, we analyze cell clusters to see whether the same cell types group together regardless of the technology used. Interestingly, we find that this overlap is limited (**Fig. 3H** and **3I**). Thus, we examined whether these subpopulations display transcriptional signatures of known cell types. Data from 10x enabled us to assign most cells to immune system categories (**Fig. 3H**). Although very similar, analysis of the Parse Evercode WT data assigned significantly more cells to unknown cell-type clusters (**Fig. 3I**). However, this analysis may introduce bias because of the batch correction of Parse samples with 10x samples, where the latter detects a higher fraction of mature mRNAs. Thus, it may mask cell-type detection by Parse, which may be less sensitive to transcriptional signatures present in the nuclei, as mature mRNA is usually assigned in this type of analysis.

These results indicate that both platforms detect similar cell types, but in different proportions, and support our earlier conclusion that the bias toward polyadenylated transcript detection in 10x may yield more physiologically relevant data. Nonetheless, Parse may still detect non-polyadenylated transcripts and provide insights into nascent transcripts, transcription of non- coding regions, or regulatory elements. Finally, data from both platforms support the idea of the heterogeneous nature of GM12878 LCLs (41).

### Transcriptional heterogeneity is reproducible across platforms and varies over the cell cycle

An important use case of single-cell transcriptomics is to quantify variation in individual gene expression across a cell population. In most scRNA-seq experiments, cells come from a heterogeneous population, and differences in gene expression levels predominantly arise because cells belong to different cell types, enabling cell-type classification. In our experiments, we assumed the cells were homotypic (lymphoblastoid cell line GM12878), yet we still observed variation in expression; we define this variation as transcriptional heterogeneity, also known in the literature as transcriptional noise (42,43).

To quantify transcriptional noise, we noted that raw variance in gene UMI counts was uninformative, since variance was strongly coupled to expression level. As shown in **Figure 4A**, the variance of log-normalized UMI counts rose steeply with the mean before saturating, a signature of the sampling statistics inherent to counting rare events. We therefore considered the residual dispersion, obtained after detrending this mean–variance relation, as our measure of transcriptional noise (**Fig. 4B**). Two complementary approaches were used to obtain this quantity: a non-parametric stratified *z*-score and a model-based estimate from the package *Bayesian Analysis of Single-Cell Sequencing* (*BASiCS*) (44), where we used the fitted residual overdispersion parameter *ɛ*. Both gave concordant results (**Supplementary Fig. S4**), and full details of both procedures are given in the Materials and Methods section.

**Figure 4.**
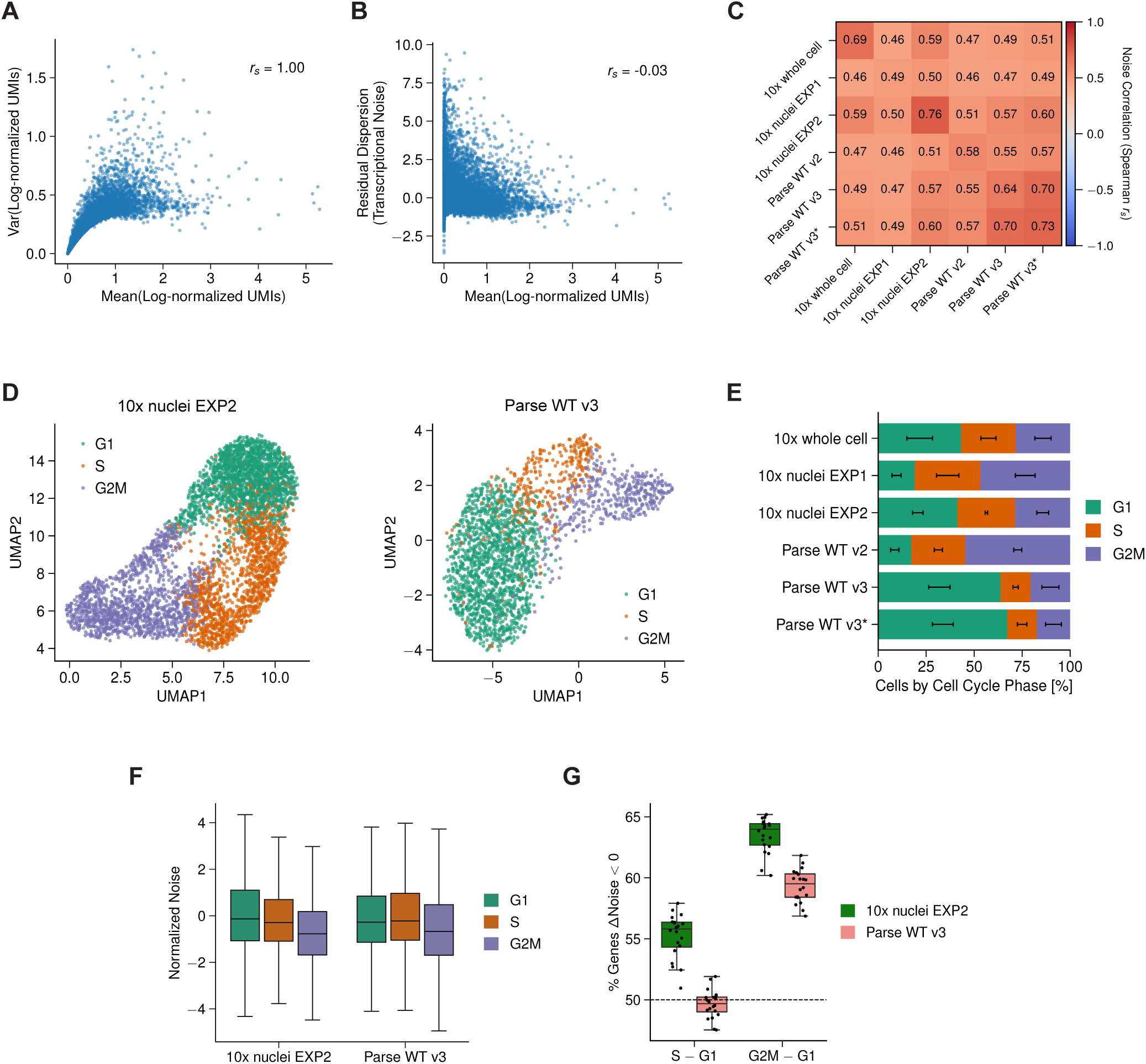
Transcriptional noise is reproducible across platforms and reduced in G2M. **(A)** Scatter plot showing the variance against mean of the log-normalized UMI counts for genes, with Spearman rank correlation reported. Data are from Replicate A of the experiment 10x nuclei EXP2. **(B)** Scatter plot showing the residual dispersion of the log-normalized UMI counts, which is our definition of transcriptional noise based on stratified *z*-score, against mean after detrending the mean–variance relation, with Spearman rank correlation reported. Data are from the same replicate in (A). **(C)** Mean Spearman rank correlation of the stratified *z*-score between experiments. Diagonal entries give the mean correlation between pairs of replicates within an experiment (excluding self-correlation); correlations were computed over the common set of genes detected in all six experiments. **(D)** UMAPs of cell-cycle marker gene expression, with cells colored by their assigned cell-cycle phase (G1, S, and G2M), for Replicate A of the experiments 10x nuclei EXP2 (left) and Parse WT v3 (right). **(E)** The relative proportion of cells in each cell-cycle phase for all experiments. Proportions were computed per replicate then averaged. The error bar within each bar reports the standard deviation of the proportion of cells in a particular phase across replicates. **(F)** Distributions of transcriptional noise for genes in each cell-cycle phase, for 10x nuclei EXP2 (left) and Parse WT v3 (right). Cells were subsampled to per phase so that all phases were based on equally many cells. Boxes show the interquartile range and median, with whiskers extending to 1.5× the interquartile range and outliers not shown. Cell-cycle marker genes and genes with zero variance in any subsample were excluded (see SI). Data are from Replicate A of each experiment. **(G)** Boxplots showing the proportion of genes with reduced noise in S or G2M relative to G1, over 20 independent subsamples of cells per phase. Each point is one subsample, and boxes show the interquartile range and median, with whiskers extending to 1.5× the interquartile range. The dashed line marks the 50% expected under no difference. Genes with identical estimates in both phases were counted in neither direction.

To assess whether this metric captured a reproducible property of each gene rather than dataset-specific fluctuation, we compared per-gene noise scores between experiments, for the common set of genes across all six experiments (**Fig. 4C**). Spearman rank correlations between replicates of the same experiment reached *r_s_*= 0.49–0.76, while correlations between independent experiments were only modestly lower (*r_s_* = 0.46–0.70), indicating that the same genes were consistently identified as more or less noisy across independent samples and across capture chemistries.

Because the cells in these datasets were unsynchronized, cell-cycle phase could have contributed to the variation captured by our noise metric. We therefore assigned each cell to G1, S, or G2M based on the relative expression of phase-specific marker genes (see Materials and Methods). The resulting assignments separated cleanly in a two-dimensional embedding of marker-gene expression (**Fig. 4D**). The proportion of cells in each phase varied considerably between experiments (**Fig. 4E** and **Supplementary Fig. S5**), with G1 ranging from 14.5% to 71.7%. This variation may reflect undetected microenvironmental fluctuations during cell culture or platform-specific technical artifacts.

To assess whether noise varies across cell-cycle phases, we randomly subsampled the data so that each phase contributed the same number of cells (*n* = 300, set by the smallest phase across datasets). This ensured that any differences between phases did not simply reflect one phase being measured in more cells than another. The noise metric showed that genes were consistently less variable in G2M than in G1 (**Fig. 4F**). We quantified this shift as the proportion of genes with reduced noise in the later phase, computed over 20 subsamples per replicate (**Fig. 4G**; see Materials and Methods). The median of this proportion was 64.0% (range 60.2–65.2%) in 10x nuclei EXP2 and 59.5% (range 56.9–61.8%) in Parse WT v3, both consistently above the 50% expected under no change. This shift was likely constrained by the number of cells available, since per-gene dispersion estimates are less precise at smaller sample sizes, which attenuates any true difference: repeating the analysis in 10x nuclei EXP2 at the larger sample size permitted by that dataset (*n* = 1,100 per phase) raised the proportion to 65.7–70.6% for G2M (ranges across the four replicates; **Supplementary Fig. S5B** and **S5C**).

Comparing S with G1, the trend was less clear, and the two experiments disagreed. In 10x nuclei EXP2, the proportion was 55.8% (51.0–57.9%), above 50% in every subsample, and increased to 58.0–61.1% at the larger sample size; in Parse WT v3, it was 49.7% (47.5–51.9%), spanning the null. S-phase cells carry a mixture of replicated and unreplicated loci and are the least reliably assigned of the three phases.

### Transcriptional noise is associated with gene length and promoter architecture

Having established that our noise metric was reproducible across experiments, we asked which gene properties it was associated with. Because gene properties often vary systematically with expression level, we assessed each association separately within low-, mid-, and high-expression strata. The most prominent association we found was with gene length: genes with larger genomic spans were consistently more likely to be among the noisiest genes. (**Fig. 5A**), an effect present at all three expression levels and in every experiment examined (**Fig. 5B**; *r_s_* = 0.20–0.47 for low, 0.46–0.59 for mid, and 0.30–0.42 for high expression). The association was weakest in the low-expression stratum, where per-gene dispersion estimates were least precise and any true correlation was attenuated. Because the same trend emerged in both 10x and Parse experiments, and in both whole-cell and nuclear preparations (see **Supplementary Fig. S6 and S7**), we conclude that this relationship represents a fundamental biological feature of gene regulation rather than a chemistry-specific artifact.

**Figure 5.**
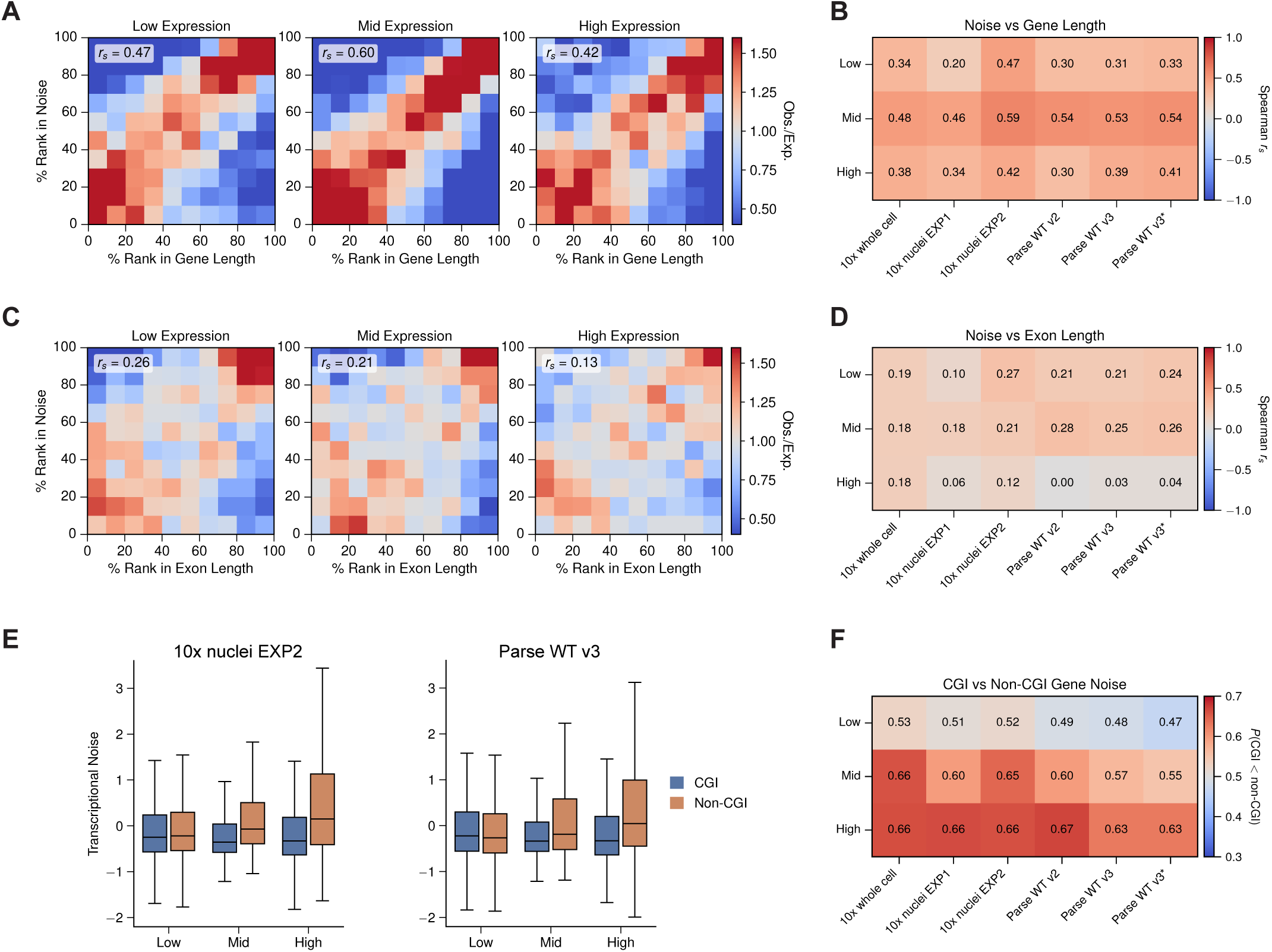
Gene-intrinsic properties associated with transcriptional noise. **(A)** Joint distribution of transcriptional noise and gene length, shown separately for genes in the low, mid, and high level of mean expression. Genes were divided into deciles along each axis, and the color of each bin gives the observed number of genes relative to that expected under independence. Spearman rank correlation between noise and gene length is shown for each expression stratum. Gene length is the genomic span from transcription start to end. Data are from Replicate A of 10x nuclei EXP2, and equivalent panels for 10x whole cell and Parse WT v3 are shown in **Supplementary Figures S6A-C**. **(B)** Mean Spearman rank correlation between noise and gene length for each experiment and expression stratum, averaged across replicates. Values for individual replicates are shown in **Supplementary Figure S6D**. **(C)** As in (A), using total exon length in place of genomic span. Total exon length is the summed length of the merged exons of a gene, corresponding to the mature transcript. **(D)** As in (B), for total exon length. **(E)** Distributions of transcriptional noise for genes with and without a CpG island promoter, by expression stratum, for 10x nuclei EXP2 (left) and Parse WT v3 (right). Genes were classified as CGI genes if their transcription start site fell within 1 kbp of an annotated CpG island (see SI). Boxes show the interquartile range and median, with whiskers extending to 1.5× the interquartile range and outliers not shown. Data are from Replicate A of each experiment. **(F)** Probability that a randomly chosen CGI gene has lower noise than a randomly chosen non-CGI gene, for each experiment and expression stratum, averaged across replicates. Values above 0.5 indicate reduced noise in CGI genes. Values for individual replicates are shown in **Supplementary Figure S8**.

To dissect the molecular basis of this association, we repeated the analysis using total exon length rather than genomic span. The correlation was substantially weaker across all expression levels (**Fig. 5C** and **D**), and in the high-expression stratum, where dispersion was most reliably estimated, it was essentially absent in several experiments (*r*_s_ 0.00–0.18). Because a 3′-capture assay counts mature transcripts, the correlation is unlikely to reflect effects such as capture efficiency or transcript-length-dependent UMI recovery. Instead, the association reflects the genomic extent over which a gene is transcribed. We note that this cannot be attributed to nuclear pre-mRNA content alone, as the correlation is equally present in whole-cell preparations, where the RNA pool is predominantly spliced (**Supplementary Fig. S6** and **S7**).

Finally, we examined the relation between transcriptional noise and promoter architecture. We found that genes whose transcription start site overlaps a CpG island were less noisy than those without, in both datasets and at mid and high expression levels (**Fig. 5E, 5F** and **Supplementary Fig. S8**). Expressing this shift as the probability that a randomly chosen CGI gene is less noisy than a randomly chosen non-CGI gene, we obtained 0.55–0.66 at mid expression and 0.63–0.67 at high expression across all six experiments, compared with 0.47– 0.53 at low expression, where the metric is least reliable.

## Discussion

This study compared two widely used single-cell transcriptomic platforms, 10x Chromium and Parse Evercode WT, using the isogenic GM12878 human lymphoblastoid cell line. Because these platforms rely on fundamentally different barcoding and reverse-transcription chemistries, they provide a useful opportunity to separate platform-dependent technical effects from biological variation. Our work extends previous comparative studies by focusing specifically on nuclear transcript detection and by asking whether transcriptional heterogeneity, or cell-to-cell variation in gene expression, can be measured consistently across assays.

The clearest practical difference between the platforms was how reliably each delivered the intended experiment. Parse Evercode WT achieved nearly 100% of the targeted sequencing depth and nuclei count, consistently across replicates and at two target depths. By contrast, 10x Chromium missed its target depth in every experiment, overshooting by approximately twofold in two cases and undershooting in a third. This difference likely reflects when cells can be counted in each workflow: Parse allows counting immediately before lysis, whereas 10x allows counting only before cells are loaded into the microfluidic device. The strong technical reproducibility of Parse has been reported previously (29), although with fewer replicates; our four-replicate design confirms this observation and points to its likely cause. Conversely, the higher cell retention observed for 10x is consistent with earlier comparisons across cell lines (30). However, overloading has a clear cost, as shown by the elevated doublet rate in 10x nuclei EXP2, where doublets reached approximately 15%, compared with 8–10% in the other experiments.

Despite differences in experimental predictability, the two platforms performed similarly in sequencing efficiency. Although the Parse barcode is longer than the 10x barcode (48 bp versus 28 bp), differences in UMI and poly(dT) length offset this difference, yielding comparable usable-read fractions. Both platforms detected approximately 3,500 genes per nucleus at 50,000 reads, and our experiments showed that 100,000–150,000 read pairs per nucleus is near sequencing saturation. This depth is well above the 20,000 reads per cell recommended by either manufacturer for whole-cell experiments, and no nucleus-specific guidance is currently available.

The primary biological difference between the platforms lies in reverse-transcription chemistry. Oligo(dT) priming in 3′ 10x restricts capture to polyadenylated transcripts and produces a pronounced 3′ bias. In contrast, Parse’s use of random hexamers and oligo(dT) primers yields more uniform gene-body coverage, a higher intronic read fraction, and a clear preference for longer exons and longer genes, consistent with previous benchmarking studies. These biases are not global; rather, they affect specific gene classes and often act in opposite directions. Parse’s intronic capture enables detection of nascent and non-polyadenylated transcription that oligo(dT)-based capture cannot reach. For example, *TALAM1*, which lacks a poly(A) tail, was detected almost exclusively by Parse. Conversely, 10x detected polyadenylated species and canonical LCL markers more sensitively, including *SNHG3*, *IGHD*, *CD79A*, and *CD81*. A study using only one of these platforms could therefore reach opposite conclusions about whether *TALAM1* or *SNHG3* is expressed in GM12878. Ribosomal protein-coding genes remain an unexplained exception, as they were better detected by 10x despite being polyadenylated and, in principle, accessible to both strategies. This observation reproduces earlier reports (30), but, as in those studies, it is not explained by GC content, which did not differ between platforms in our data.

Apart from these platform-specific biases, agreement on gene-level mean expression was strong across platforms and replicates. This suggests that the chemistries differ mainly in which transcript classes they capture, rather than in how much of a given gene they detect once that gene is captured. This distinction matters for reusing aggregated resources such as CellxGene curated by the Chan Zuckerberg Initiative (45) and singleCellBase (46). In our data, cells clustered by platform rather than by biological state, and *Harmony* did not integrate them sufficiently to compare cell-type assignments on equal footing. This failure in data integration might lead to errors in cell-type assignment. Similar masking has been reported across preparations and platforms, highlighting the need for deliberate calibration of correction methods (47). We therefore suggest that integrations spanning distinct barcoding chemistries should either be restricted to, or validated on, the polyadenylated genes common to both approaches.

Quantifying transcriptional noise requires careful correction for mean expression. Raw variance in UMI counts is dominated by mean expression, a long-recognized problem in noise analysis: coefficient-of-variation measures are negatively correlated with expression level, which causes unadjusted metrics to systematically identify lowly expressed genes as the noisiest (48). We therefore detrended the mean–variance relationship and used residual dispersion as our measure of noise, calculated independently using a stratified *z*-score and the BASiCS overdispersion parameter (44). Because different dispersion estimators can disagree substantially on which genes are noisy (49,50), we report concordance between two independent estimators.

A central question is whether residual dispersion reflects an intrinsic property of the gene or an assay artifact, since dispersion can be affected by capture efficiency and detection bias. Per-gene noise ranks were concordant across independent experiments (*r_s_* = 0.46–0.70), only modestly lower than the concordance observed between replicates of the same experiment (*r_s_* = 0.49–0.76), and this pattern held for both platforms. An artifact specific to oligo(dT) capture should not persist in random-hexamer data, and the reverse should also be true. To our knowledge, this is the first demonstration that a detrended noise metric is transferable between fundamentally different barcoding chemistries. It therefore provides an empirical basis for comparing noise estimates across the increasingly platform-heterogeneous public datasets.

Cell-cycle dependence has previously been shown to dominate expression noise in yeast (51), and mammalian cells partially compensate for gene dosage through cell-size-dependent transcription (52). In our study, genes were consistently less variable in G2M than in G1 across both platforms. Gene dosage offers a straightforward explanation: after replication, transcription proceeds from four rather than two independently bursting templates and averaging over more copies reduces relative variance. This model predicts a gradient based on replication timing within S phase, which may explain why our S–G1 comparison was equivocal and differed between platforms. However, a technical explanation remains possible. G2M cells are larger and contain more RNA, and higher counts may deflate dispersion in ways that normalization does not fully remove. We therefore regard the dosage interpretation as provisional pending count-matched downsampling.

Among the gene attributes examined, gene length was the strongest correlate of noise, whereas the association with total exonic length was substantially weaker and essentially absent at high expression in several experiments. This dissociation has a direct precedent: Larsson *et al.* reported that gene length, but not spliced mRNA length, inversely correlated with burst size, with no effect on burst frequency (53). The fact that two separate methods-allele-resolved burst inference and detrended dispersion- both differentiate gene length from exonic length indicates that this effect is real and mechanistically connected to intronic sequences, rather than being influenced by transcript size or capture.

In a two-state model, smaller bursts imply lower dispersion, so the Larsson *et al.* result would predict that longer genes are less noisy (53), whereas we observe the opposite. We see three non-exclusive explanations. First, the measurements differ by compartment: allele-resolved burst kinetics were derived from mature cytoplasmic mRNA, whereas our nuclear measurements retain intronic and nascent signal, and nuclear retention is known to buffer cytoplasmic variability (54). In addition, nuclear noise has been reported to be higher than cytoplasmic noise (55). Second, the telegraph model does not explicitly represent elongation; a long genomic span delays transcription initiation and transcript completion, adding a source of variance not captured by burst size or frequency. Third, introns contain regulatory elements, pausing sites, and opportunities for premature termination, each of which can introduce stochastic steps downstream of initiation. We also note that not all studies recover a gene-length association; one recent assessment found the relationship weak and non-significant (49). In this context, the consistency of our effect across two scRNA-seq platforms, three expression strata, and nuclear and whole-cell preparations provides the strongest evidence for its robustness.

Another possibility is that gene length affects promoter architecture. Longer templates generate more torsional stress during elongation, and if topoisomerase-mediated relaxation is slow, supercoiling buildup could increase the time between successful initiation events. This mechanism is established in bacteria, where gyrase inhibition increases bursting (56), and it has been modeled for eukaryotic divergent transcription (57). Consistent with this possibility, genes with CpG-island transcription start sites were less noisy than non-CGI genes at mid and high expression levels across all experiments presented here. This reproduces an established relationship in which CGI presence lowers expression variability while TATA boxes increase it (58), but does so using a metric and platform pairing not previously applied to this question. CGI promoters are predominantly bidirectional, and divergent initiation deposits negative supercoiling into the shared promoter region, a configuration that favors duplex opening. Their constitutive nucleosome-depleted regions may also reduce the slow chromatin-remodeling transitions that generate long off-states at other promoters. Together, these features could buffer promoter activity both torsionally and kinetically.

Overall, resolving transcriptional heterogeneity and noise using single-cell transcriptomic approaches appears feasible, though important challenges remain. Parse Evercode WT provides stronger technical reproducibility, more predictable recovery of cell number and consistent sequencing depth than 10x Chromium. At the same time, biological contributors to transcriptional variation must be accounted for. In tissues, cell-type composition substantially contributes to cell-to-cell heterogeneity; in a comparatively homogeneous cell line such as GM12878, our results show that cell-cycle phase is a major contributor. The main limitation of our approach was using unsynchronized cells. For 10x Chromium experiments, we recommend shallow sequencing to establish the recovered cell number before adjusting the final sequencing depth. In Parse experiments, by contrast, cell number is more predictable, allowing accurate sequencing depth to be targeted from the outset.

## Resource availability

### Lead Contact

Requests for further detailed information and resources should be directed to the lead contact, Prof. Nick Gilbert, and co-corresponding author Dr Rafal Czapiewski.

### Materials availability

This study did not generate new, unique reagents.

### Data and code availability

All raw data will be available from public depositories upon acceptance of this study. Supplementary data are available in a separate file with the submission. All the data and code needed to reproduce the figures in this manuscript will be available from public depositories upon acceptance of publication. We list open-source software pipelines in the methods section.

## Acknowledgements

We thank Susan Campbell, Michael Rennie, and Elizabeth Freyer from the 10x facility at the MRC Human Genetics Unit, University of Edinburgh, for their support. We also thank Dr. Meryam Beniazza from the Institute for Regeneration and Repair, Single Cell & Spatial Biology Facility at the University of Edinburgh, for assistance with the 10x Chromium single-nucleus experiment. We appreciate the members of the Gilbert Lab and Marenduzzo Lab for their discussions and comments on our manuscript. This work was funded by grants from the Wellcome Trust (223097/Z/21) and the UK Medical Research Council (MC_UU_00007/13), both awarded to Nick Gilbert.

## Author contribution

Study Conceptualization, R. C, M. C., D. M. and N. G.; methodology, R. C., C. N., and N. G.; software, M. C., J. D. and G. G.; formal analysis, R. C., M. C., G. G. and J. D.; validation, R. C. and M. C.; investigation, R. C. and C. N.; data curation, R. C., M. C. and G. G.; writing – original draft, R. C., M. C., J. D., D. M. and N. G.; writing – reviewing & editing, R. C., M. C., J. D. and N. G.; visualization, R. C., M. C. and J.D.; supervision, N. G. and D. M.; project administration, R. C. and N. G.; Funding acquisition, N. G.

## Declaration of generative AI and AI-assisted technologies

No large language models (LLMs) were used to assist with the manuscript writing process. The *Microsoft Word* built-in *Editor* function was used to check spelling and grammar. Furthermore, *Grammarly Pro* software was used to correct spelling, grammar, and readability. No generative AI technologies were used to prepare the text, figures, and tables.

## Supplementary Information

Document S1: Table S1 and Figures S1-S8 as one PDF.

## Materials and Methods

### GM12878 cell culture and technical replicates

The Lymphoblastoid GM12878 cell line was cultured in RPMI-1640 media supplemented with 15% fetal bovine serum, penicillin (100 U/mL), and streptomycin (100 μg/mL). Cells were routinely seeded at a density of 0.2 × 10^6^/mL and used in experiments or split at 1.8-2.0 × 10^6^/mL. Routine cell culture was maintained in 20 mL of media in a T-75 flask at 37°C in a 5% CO_2_ atmosphere. Cells were regularly tested for mycoplasma. The GM12878 population doubling time is reported to be approximately 24 hours, and we observed similar results. Growth was typically arrested at a concentration of 1.5 × 10^6^/mL, and we observed increased cell death at this point.

Two days before nuclei isolation, cells were split into four flasks, which were treated as four separate technical replicates. Single-cell studies are often conducted with duplicates, which are insufficient for robust statistical analysis. This is usually due to financial constraints rather than deliberate choices based on statistical power calculations. Some sources recommend at least three replicates for scRNA-seq, as this enables statistical testing and outlier detection. Ultimately, triplicates provide a more dependable estimate of variability compared to duplicates, and increasing the number of replicates from two to three typically significantly enhances precision and the ability to estimate variability (59–61). Moving from three to four replicates can still improve precision and statistical power, but the benefits are smaller than increasing from two to three. The reduction in SEM when adding a fourth replicate is approximately 13%, and including a fifth replicate offers an additional 10% decrease in SEM, which may not be cost-effective. Therefore, we decided to perform our experiments with four replicates.

The cell number used in our experiments was chosen arbitrarily at the maximum recommended by the manufacturers. However, for predicting cell numbers, a prediction tool is recommended, e.g., *How Many Cells* from the Rahul Satija lab (https://satijalab.org/howmanycells/). Detailed numbers of targeted and yielded cells in each experiment and sample are listed in **Table 1** and **Supplementary Table S1**.

### GM12878 cell nuclei isolation

To obtain nuclei for experiments, we used a publicly available protocol from 10x Genomics (10x Demonstrated Protocol, CG000365). The same nuclei isolation protocol was employed for experiments on the 10x Chromium and Parse WT Evercode (SPLiT-seq) platform. Each sample in all single-nucleus experiments represents a separate T-75 culture flask, and cells from each flask were used in separate nuclei isolation preparations (**Fig. 1**). Two million GM12878 cells per sample were utilized for the nuclei isolation protocol. First, 2 × 10^6^ cells in suspension were centrifuged at 300x *g* for 5 minutes at 4 °C to remove media (a swing-bucket centrifuge is recommended); next, the supernatant was discarded, and the cell pellet was washed once with PBS supplemented with 0.04% BSA at 4 °C. After the wash, cells were centrifuged at 300x *g* for 5 minutes at 4 °C and resuspended in 100 μL of chilled Lysis Buffer [10 mM Tris-HCl pH 7.4, 10 mM NaCl, 3 mM MgCl_2_, 1% BSA, 0.1% Tween-20, 0.1% NP-40, 1% BSA, 1 mM DTT and 1 U/μL RNase inhibitor (Roche cat: 3335402001)] and incubated on ice for 5 minutes. Immediately after lysis, 1 ml of Wash Buffer was added (10 mM Tris-HCl pH 7.4, 10 mM NaCl, 3 mM MgCl_2_, 1% BSA, 0.1% Tween-20, 1 mM DTT, and 1 U/μL RNase inhibitor). After gently pipetting up and down 5 times, the mixture was centrifuged at 500x *g* for 5 minutes at 4 °C, and the supernatant was discarded. Then, 1 mL of Wash Buffer was added to the pelleted nuclei, which were resuspended and then centrifuged at 500x *g* for 5 minutes at 4 °C. These steps were repeated for a total of three washes. Finally, the number of nuclei was determined by a Neubauer hemocytometer. Typically, we obtained approximately 1 million purified nuclei from 2 million cells in the input material. The cell aggregation, integrity of the nuclei, and the purity of the preparation were confirmed by analyzing images of DAPI-stained nuclei and images from a phase contrast microscope (**Supplementary Fig. S1C** and **S1D**).

### 10x Chromium cDNA library preparation and sequencing strategy

Freshly isolated nuclei were used immediately after counting in the 10x Chromium protocol, following the manufacturer’s recommended concentrations (10x protocol, CG000338). For the 10x whole cell experiment, cells were washed in PBS and sorted via FACS to remove aggregates. Next, 16,000 cells per replicate (6000 cells/μL) were loaded into the experiment. For the 10x nuclei EXP1, 12,600 nuclei per replicate (5000 nuclei/μL), and 17,000 nuclei per replicate (4900 nuclei/μL) for the 10x nuclei EXP2. We used the Chromium Next GEM Single Cell Multiome ATAC + Gene Expression Reagent Bundle (cat. 1000285) and Chromium Next GEM Chip J (cat. 1000230), following the manufacturer instructions. Representative TapeStation traces of QC profiles for cDNA and sequencing libraries are shown in **Supplementary Figures S1E** and **S1F**.

The recommended sequencing cycle numbers are: 28 cycles for read 1, covering the 10x barcode and UMI barcode; 10 cycles for i7 index; 10 cycles for i5 index; and at least 90 cycles of read 2 for the cDNA insert (**Supplementary Fig. S1G**). Additionally, the samples were spiked with 1% PhiX libraries (**Figure S1** and **Table S1**). For all experiments, sequencing was outsourced to external vendors listed in Supplementary Table S1. The Illumina chemistry version and the actual cycle numbers used for sequencing each experiment are also provided in Table S1.

### Parse SPLiT-seq cDNA library preparation and sequencing strategy

Immediately after nuclei isolation, one million freshly isolated nuclei per sample were processed using the *Parse Evercode Fixation* protocol version 2.1.1, following the manufacturer’s instructions. Briefly, nuclei were permeabilized and fixed in proprietary PFA-based solutions, then 1 × 10^6^ cells per sample were frozen in a container at −80 °C in a proprietary DMSO-based solution. To assess consistency, reproducibility, and the manufacturer’s claims of improved cell retention and transcript detection sensitivity, we performed Parse SPLiT-seq experiments using *Parse Evercode WT Mini* kits version 2 and 3, which provided manual versions 1.2 and 1.3, respectively. In both experiments, we used the option with Unique Dual Indices (UDIs), ensuring the libraries contain 5’ and 3’ TruSeq adapters. After thawing fixed cells and recounting them, we did not observe significant cell loss. Following the manufacturer recommendations, we used samples with less than 5% nuclei in aggregates to prevent sequencing doublets (**Supplementary Fig. S1D**). Evercode WT Mini kits are designed for 10,000 cells. However, the manufacturer states that up to 20,000 cells can be detected and divided into up to 12 samples. To reach these numbers, we used the Parse cell calculations tool and loaded 27541 cells per sample (**Fig. 1D** and **Supplementary Table S1**).

The Parse protocol involves three rounds of split-pool barcoding of cells in 96-well plates, during which mRNA is labeled in intact cells or nuclei. The fourth round of barcoding is done after cells are counted and divided into sub-libraries (**Supplementary Fig. S1A**). In our setup, cells were split into two equal sub-libraries of 10,000 cells. The Parse manual suggests sequencing the libraries with at least 64 cycles for read 1 to cover the cDNA insert, 8 cycles for the Illumina index i7, 8 cycles for i5, and 58 cycles for read 2 to cover Parse barcodes BC1-BC3. These are minimum recommended values, and the manufacturer advises increasing them if Illumina chemistry permits. For quality control, the samples were spiked with 5% PhiX libraries as advised by the manufacturer (**Fig. 1B** and **Supplementary Fig. S1G**).

### Sequencing depth

As discussed in the results section, manufacturers of both platforms recommend a minimum sequencing depth of 20,000 paired-end reads per cell. We assumed that for nuclei, which have fewer transcripts available than whole cells, we would sequence single-nucleus experiments at 50,000 reads per nucleus. Additionally, we conducted one experiment for each platform, aiming for 150,000 reads per nucleus. For our 10x whole-cell experiment, we targeted 25,000 reads per cell (**Supplementary Table S1**).

### Data analysis

The manufacturer-recommended software was used to process FASTQ files from the sequencing to obtain gene expression matrices. For the 10x Chromium, *Cell Ranger* was used; for Parse SPLiT-seq, *Split-Pipe* was used, both available from the websites of the manufacturers. Both companies also offer free web-based platforms to analyze the data using the same pipelines: *Trailmaker* from Parse and the *10x Genomics Cloud*. Many studies systematically compared these analysis pipelines on different cell lines, cell types, and dissociated tissues. These studies concluded that *STAR-solo* and *Split-Pipe* are the most refined and efficient pipelines to analyze SPLiT-seq data. Both methods produce very similar results (30,62,63). A recent review examined dimension reduction and compared methods used to reduce the dimensionality of scRNA-seq data (64). In this study, the *Harmony* method was used to remove batch effects and reduce dimensionality across experiments and technical replicates (40). Sequences were aligned to the hg38 reference genome. *RSeQC* v5.0.4 (65) was used to calculate gene body coverage, using reads that overlapped a set of ∼ 3800 housekeeping genes, as defined by Eisenberg and Levanon (66). *RSeQC* was also used to assess the genomic distribution of reads, using GENCODE (v48) as the reference gene model.

### Gene annotation

We considered different gene annotations and categories when comparing the performance of Parse and 10x Chromium technologies. To define gene categories, we downloaded the primary assembly GTF file from GENCODE (version 48), which contains the gene annotation list (67). We used the *gene_type* field to identify protein-coding and long non-coding (lnc)RNA genes. Ribosomal protein-coding genes were identified by the *RPS* and *RPL* prefixes, while mitochondrial genes were identified based on the prefix *MT* in the HGNC gene symbols. Transcription factor (TF) genes were determined using the list downloaded from the database *AnimalTFDB* (v4) (68). We also utilized the GTF file to compute gene length and total exon length. The gene length was defined as the difference between the *end* and *start* fields, and exon length was taken to be the non-overlapping union of all exonic regions of the gene. The GC content of each gene was obtained from Ensembl BioMart (v114). When examining the transcriptional heterogeneity (noise) of genes, we also studied its difference between genes whose promoters are near a CpG island (CGI genes) and those that are not (non-CGI genes). To identify CGI-associated genes, CpG islands were retrieved from the UCSC Genome Browser (hg38 assembly). We obtained the transcription start site (TSS) of genes from the MANE Select transcript set (69), which provides a single representative transcript per protein-coding gene agreed between RefSeq and Ensembl. For genes without a MANE Select transcript, predominantly non-coding genes, the strand-aware Ensembl gene start coordinate was used instead. We then classified CGI genes as those whose promoter region (TSS ± 1 kbp) intersects at least one CpG island, computed using *bedtools intersect*.

### Doublet detection

A technical artifact in single-cell transcriptomics is the presence of doublets – cells sharing the same set of barcodes. The R software package *scDblFinder* was used to detect and remove these cells using the default parameters (32). Specifically, count matrix data (without filtering) from *Split-Pipe* and *Cell Ranger* were loaded as a *SingleCellExperiment* object and passed to the function *scDblFinder*, which then classified a cell as a singlet or doublet.

### Filtering and normalization

Count matrices produced by *Split-Pipe* and *Cell Ranger* were filtered to remove cells and genes with low counts. Cells with less than 1,500 UMI counts were removed, as well as cells having greater than 20% of the UMIs from mitochondrial genes or being classified as doublets by *scDblFinder*. For analysis of transcriptional heterogeneity, a more stringent level of filtering was performed to ensure that the mean–variance relation was stable for normalization (see below). Here, we retained only cells detecting between 200 and 8,000 unique genes and with at least 10,000 UMIs. For genes, we kept only those that have counts in at least 3 cells.

After filtering, UMI counts were normalized using the Python package *scanpy* according to standard practice in single-cell transcriptomics. Specifically, UMI counts per cell were rebalanced using the method *pp.normalized_total* so that all cells have a similar number of counts. Letting *m_ij_* be the UMI count for gene *j* in cell *i*, we performed the transformation

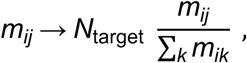

where *N*_target_ is the target total UMI counts for a cell, equal to the median of total counts for cells before normalization (this is the default option of the method). Because UMI counts span several orders of magnitude across genes, we additionally applied a log1p transformation [i.e., *m_ij_* → ln (1+*m_ij_*)] to prevent highly expressed genes from dominating downstream analyses. Unless specified otherwise, all subsequent analyses were conducted based on this transformed count matrix. Additionally, to enable comparison between experiments from different platforms, we considered only the common set of genes that were detected across all experiments when performing correlation analyses.

### Cell cycle analysis

To classify the cell cycle phase of each cell, we examined the relative expression of marker genes for S and G2M phases in the cell, following the approach done in (70). The list of marker genes was obtained from (71). The cell cycle score was computed using *scanpy*’s method *tl.score_genes_cell_cycle* using the default parameters (70). Here, the mean expression of the S and G2M marker gene sets was compared to that of control gene sets sampled from the same expression bins, and cells with a positive S or G2M score were assigned to the corresponding phase, while cells scoring negative for both assigned to G1. For the cluster maps shown in **Figure 4D**, we constructed them by performing Uniform Manifold Approximation and Projection (UMAP) on the expression data for S and G2M marker genes in each cell. We first built the nearest neighbors distance matrix and a neighborhood graph using the method *pp.neighbors* with the default parameters, where distances were computed for the 15 nearest neighbors (the default value of the method). Then, we used *tl.umap* to perform the projection and show the two-dimensional embedding.

### Calculating transcriptional heterogeneity

An important part of our study was to quantify the transcriptional heterogeneity (noise) of each gene, or its variation in expression across cells. In **Figure 4A**, we showed that the raw variance of UMI counts across cells for each gene is highly correlated with its mean UMI counts, which is typical in processes involving detection of rare events (e.g., the transcripts from individual genes). Hence, a direct comparison of raw variance between genes is uninformative, as it would yield results similar to comparing expression levels.

Here, we instead used residual dispersion as our metric for transcriptional noise, which quantifies the variation in UMI counts after detrending the mean–variance relation. We employed two complementary approaches to determine this quantity: a non-parametric and a model-based method. For the former, we divided genes into 100 strata, each containing approximately an equal number of genes. Within each stratum, we then computed the *z*-score of the sample standard deviation (SD) of UMI counts for each gene. More precisely, say gene *j* belongs to stratum *s* and its SD of UMI counts is SD*_j_*, its *z*-score is

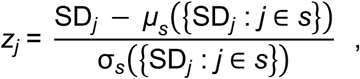

where *μ_s_* and σ*_s_* are the mean and standard deviation of the SD of UMI counts for all genes in stratum *s*. By construction, this procedure removes the systematic dependence of the noise metric on mean expression (**Fig. 4B**; *r_s_* = −0.03), in contrast to raw variance, which correlates strongly with mean UMI counts at *r_s_* = 1.00 (**Fig. 4A**).

For the model-based approach, we employed the R package *Bayesian Analysis of Single-Cell Sequencing* (*BASiCS*) to detrend the mean–variance relation (44). BASiCS models raw UMI counts with a hierarchical Poisson–Gamma formulation, in which gene-specific expression is captured by a mean parameter *μ_j_* and an overdispersion parameter *δ_j_* that quantifies variability beyond that expected from Poisson sampling alone, with cell-specific normalization and technical noise terms accounting for differences in sequencing depth and capture efficiency between cells. Because *δ_j_* retains a systematic dependence on *μ_j_*, we examined the residual overdispersion *ɛ_j_*, defined as the deviation of ln *δ_j_*from a global regression trend fitted across genes, which measures transcriptional variability that is largely decoupled from expression level. As spike-ins were unavailable, we used the separate replicates from each experiment to allow the package to estimate and correct for batch effects; as a result, only one set of fitted model parameters was obtained for each experiment but not for individual replicates. We ran the Markov Chain Monte Carlo sampler from the package for 20,000 steps, of which the first 10,000 steps were for burn-in, with sampling done every 10 steps for the remaining period.

This procedure was done using the *BASiCS_MCMC* method in the package, with options WithSpikes = FALSE and Regression = TRUE, and we set PriorMu in the PriorParam option to *EmpiricalBayes*. We took the median of the sampled residual overdispersion *ɛ_j_* as our metric for noise. **Supplementary Figure S4C** shows that both the non-parametric and model-based approach yield similar results, with Spearman rank correlation score *r_s_*between *z_j_* and *ɛ_j_* from the same replicate above 0.58 in both Parse and 10x Chromium platforms. In **Supplementary Figure S7**, we showed that *ɛ_j_* correlates with gene length and to a less extent the total exon length, while in **Supplementary Figure S8B** we found that CGI genes have a lower *ɛ_j_* compared to non-CGI genes, consistent with the results for *z_j_* (**Supplementary Figures S6 and S8A**).

Because the number of genes is large and genes are not independent, hypothesis tests comparing gene-level distributions return vanishingly small *p*-values for differences of negligible size. We therefore report effect sizes, e.g., correlation coefficients, the proportion of genes with reduced noise, or the probability that a randomly chosen gene from one group has lower noise than one from the other, together with the spread across replicates or subsamples, rather than *p*-values, for all comparative analyses.

For comparing transcriptional noise between cells in different cell-cycle phases, we considered a normalized *z*-score metric. Genes were assigned to strata and the reference statistics *μ_s_* and σ*_s_* obtained as described above, using all cells. The per-gene SD was then recomputed within each phase separately and standardized against the reference statistics of the gene’s assigned stratum, so that scores from different phases share a common scale. Note that the reference terms cancel when two phases are subtracted (see **Fig. 4G**), so the per-gene difference is proportional to the difference in raw standard deviations.

Since the precision of a per-gene SD estimate depends on the number of cells available, and cell numbers differed substantially between phases (G1, *n* = 1,821 ; S, *n* = 1,388 ; G2M, *n* = 1,369 for Replicate A of 10 nuclei EXP2, while G1, *n* = 1,458; S, *n* = 329; G2M, *n* = 432 for Replicate A of Parse WT v3), all comparisons were performed on subsamples of equal size (*n* = 300 cells per phase, matched to the smallest phase across both datasets). This procedure was repeated for 20 independent trials, and we reported the median and range across trials. We noted that because subsample size approached the population size for the smaller phases, draws shared a substantial proportion of cells and the resulting spread was a lower bound on sampling variability.

At this cell number, a fraction of genes were undetected within a given subsample and had zero sample standard deviation; such genes were excluded from the comparison, as their standardized score was determined entirely by the reference statistics of their stratum rather than by any measured variability. Genes that remained undetected in a subset of draws would produce identical estimates in both phases being compared, giving a point mass at zero in the distribution of per-gene differences. We therefore quantified the shift between phases as the proportion of genes with a lower *z*-score in the later phase, which would not be affected by these ties, rather than as a median difference.

