## Supplementary_Document_S1 for "Cross-chemistry single-nucleus RNA-seq identifies gene length and CpG-island promoters as determinants of transcriptional noise"



Figure S1

Table S1

Figure S2

Figure S3

Figure S4

Figure S5

Figure S6

Figure S7

Figure S8

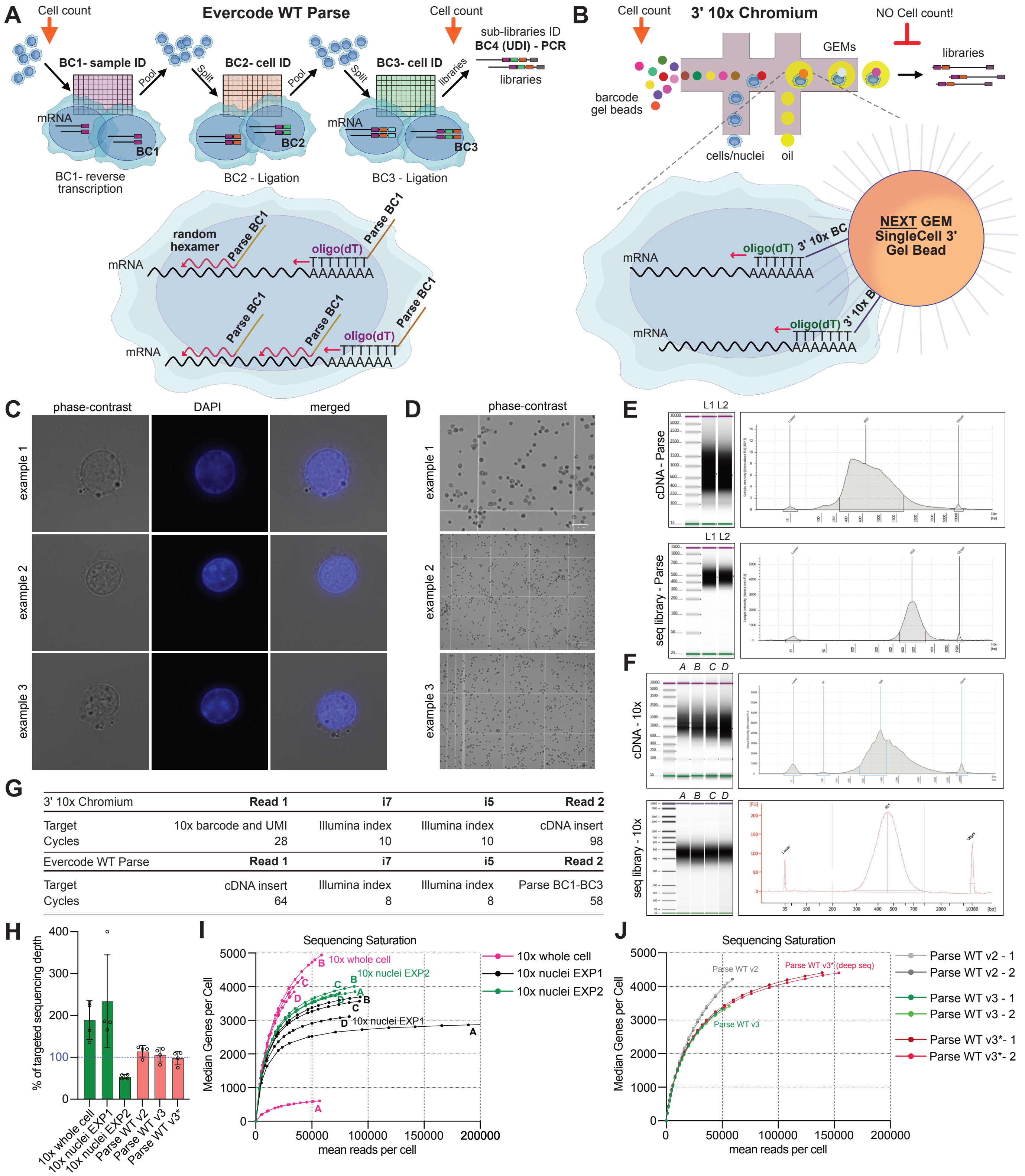


**Supplementary Figure S1: Barcoding workflow and QC comparison related to Figure 1. (A and B)** Comparison of the Evercode WT Parse and 3'10x Chromium platforms. Parse BC1, which consists of random hexamers fused with the first barcode, enables sample identification. BC2 and BC3 identify individual cells. After three rounds of barcoding, cells are counted and loaded into sub-libraries. During the PCR reaction, the final barcode—BC4—is added to identify sub-libraries. Microfluidic 10x Chromium Next GEM uses oligo(dT) fused to the 10x barcode for reverse transcription. **(C and D)** Representative images of nuclei isolated from GM12878 cells used for experiments (C), and examples of freshly isolated nuclei assessed on the Neubauer hemocytometer, imaged by the Zoe BioRad system. High-quality samples contain single, isolated nuclei with no debris or aggregates. The same protocol for nuclei isolation was used for Parse and the 10x Chromium workflow. See also the Methods section. **(E and F)** Representative traces of cDNA for both platforms (E) and traces of sequencing libraries (F). **(G)** The table with manufacturer recommendations for a dual-indexed sequencing configuration. These suggested reads can be longer; however, shorter reads may reduce transcriptome alignment rates, as noted in the manufacturer's manual. **(H)** Percentage of targeted sequencing reads yielded. **(I and J)** Sequencing saturation. Median number of genes per cell at down-sampled sequencing depths. The left panel shows four replicates for each experiment using the 10x platform (I). The right panel shows two libraries for each Parse Evercode WT experiment (J).

| Platform/Exp | **sc**RNA-seq** 10x Chromium | **sn**RNA-seq 10x Chromium EXP1 3' v3.1 | **sn**RNA-seq 10x Chromium EXP2 3' v3.1 | **sn**RNA-seq Evercode (WT) Parse v.2 | **sn**RNA-seq Evercode (WT) Parse v.3* | **sn**RNA-seq Evercode (WT) Parse v3* **deep** |
|---|---|---|---|---|---|---|
| **Number of seq libraries** | 4 | 4 | 4 | 2§ | 2§ | 2§ |
| **Number of tech. replicates** | 4 | 4 | 4 | 4 | 4 | 4 |
| **Targeted nuclei No./Exp.** | 40,000 = 10,000/libr. | 32,000 = 8,000/libr. | 40,000 = 10,000/libr. | 20,000 = 10,000/libr. | 20,000 = 10,000/libr. | 20,000 = 10,000/libr. |
| **Targeted nuclei No./sample** | 10,000** | 8,000 | 10,000 | 5,000 | 5,000 | 5,000 |
| **Loaded nuclei/ sample** | 16,000** | 12,600 | 17,000 | 27,541 | 27,541 | 27,541 |
| **Targeted reads/ nucleus** | 50k/cell | 50k/nuclei | 150k/nuclei | 50k/nuclei | 50k/nuclei | 150k/nuclei |
| **Yielded reads/ nuclei/replicate** | ***A*** - 56,000<br>***B*** - 58,000<br>***C*** - 41,000<br>***D*** - 34,000 | ***A*** - 200,000<br>***B*** - 93,000<br>***C*** - 92,000<br>***D*** - 83,000 | ***A*** - 88,000<br>***B*** - 87,000<br>***C*** - 74,000<br>***D*** - 74,000 | ***A*** - 60,000<br>***B*** - 60,000<br>***C*** - 47,000<br>***D*** - 62,000 | ***A*** - 60,000<br>***B*** - 58,000<br>***C*** - 41,000<br>***D*** - 53,000 | **A** - 166,000<br>**B** - 163,000<br>**C** - 113,000<br>**D** - 147,000 |
| **Illumina Sequencer** | NextSeq 2000 | NextSeq 2000 | NovaSeq X Plus | NovaSeq X Plus | NovaSeq X Plus | NovaSeq X Plus |
| **Read length/ cycles (PE)** | 100 cycles | 100 cycles | 150 cycles | 150 cycles | 150 cycles | 150 cycles |
| **Illumina Chemistry** | NextSeq1000/ 2000 P3 | NextSeq1000/ 2000 P3 | NovaSeq PE150 25B flow cell | NovaSeq PE150 10B flow cell | NovaSeq PE150 10B flow cell | NovaSeq PE150 25B flow cell |
| **PhiX spike** | 1% | 1% | 1% | 5% | 5% | 5% |
| **Data yield/total PE reads** | 183.3 Gb<br>1,366.8 M | 182.4 Gb<br>1,363.2 M | 1,163.9 Gb<br>3,602.3 M | 310.8 Gb<br>1,036.0 M | 384.7 Gb<br>**1,222.4 M** | 1,124.7 Gb<br>**3,352.7 M** |
| **PF clusters/ library** | ***A*** - 245,099,447<br>***B*** - 372,722,539<br>***C*** - 356,139,664<br>***D*** - 305,197,727 | ***A*** - 281,424,669<br>***B*** - 316,753,335<br>***C*** - 317,202,845<br>***D*** - 339,323,783 | ***A*** - 1,047,725,002<br>***B*** - 877,824,584<br>***C*** - 858,829,092<br>***D*** - 816,927,865 | **1** - 427,284,036<br>**2** - 608,793,177 | **1** - 561,076,716<br>**2** - 661,348,836 | **1** - 1,557,146,056<br>**2** - 1,795,580,962 |
| **Provider** | ECRF Wellcome, Edinburgh, UK | ECRF Wellcome, Edinburgh, UK | Novogene, UK | Novogene, UK | Novogene, UK | Novogene, UK |

**Supplementary Table S1. Summary of Illumina sequencing parameters related to Table 1.** Each experiment was conducted with four technical replicates (***A***, ***B***, ***C***, and ***D***). The 10x Chromium 3' platform enables the preparation of a separate cDNA library for each sample. §The Parse SPLiT-seq platform pools all samples and allows their division into sub-libraries. Our Parse setup divided samples into two sub-libraries per experiment, each containing 10,000 nuclei from all four replicates. *The last two columns on the right of the table show conditions for the same sets of libraries sequenced at different depths: **1,222.4 million** reads and **3,352.7 million** reads, assuming 50,000 and 150,000 reads per cell, respectively. **The experiment in the first column was performed using whole cells; therefore, the *cell* number is provided instead of the nuclei number. **Targeted nuclei No./Exp**—total nuclei expected from the experiment. **Targeted nuclei No./sample**—nuclei expected in each sample. **Loaded nuclei/sample**— number loaded for barcoding. **Targeted reads/nucleus** indicates the expected number of reads sequenced per cell or nucleus. **Yielded reads/nucleus/replicate** refers to the actual number of sequencing reads obtained per cell or nucleus in each replicate. The **"Illumina Sequencer"** row specifies the sequencing machine used. **"Read length/cycles (PE)"** indicates paired-end sequencing reads. **Illumina Chemistry** and **PhiX spike** describes the protocol and sequencing setup. **Data yield/total PE reads** refers to the total data output and total read number per experiment. **PF clusters/library** represents the number of raw reads yielded per library. **Provider** indicates the sequencing facility. For details on the experiments' gene and cell counts, see also the main text in Table 1.

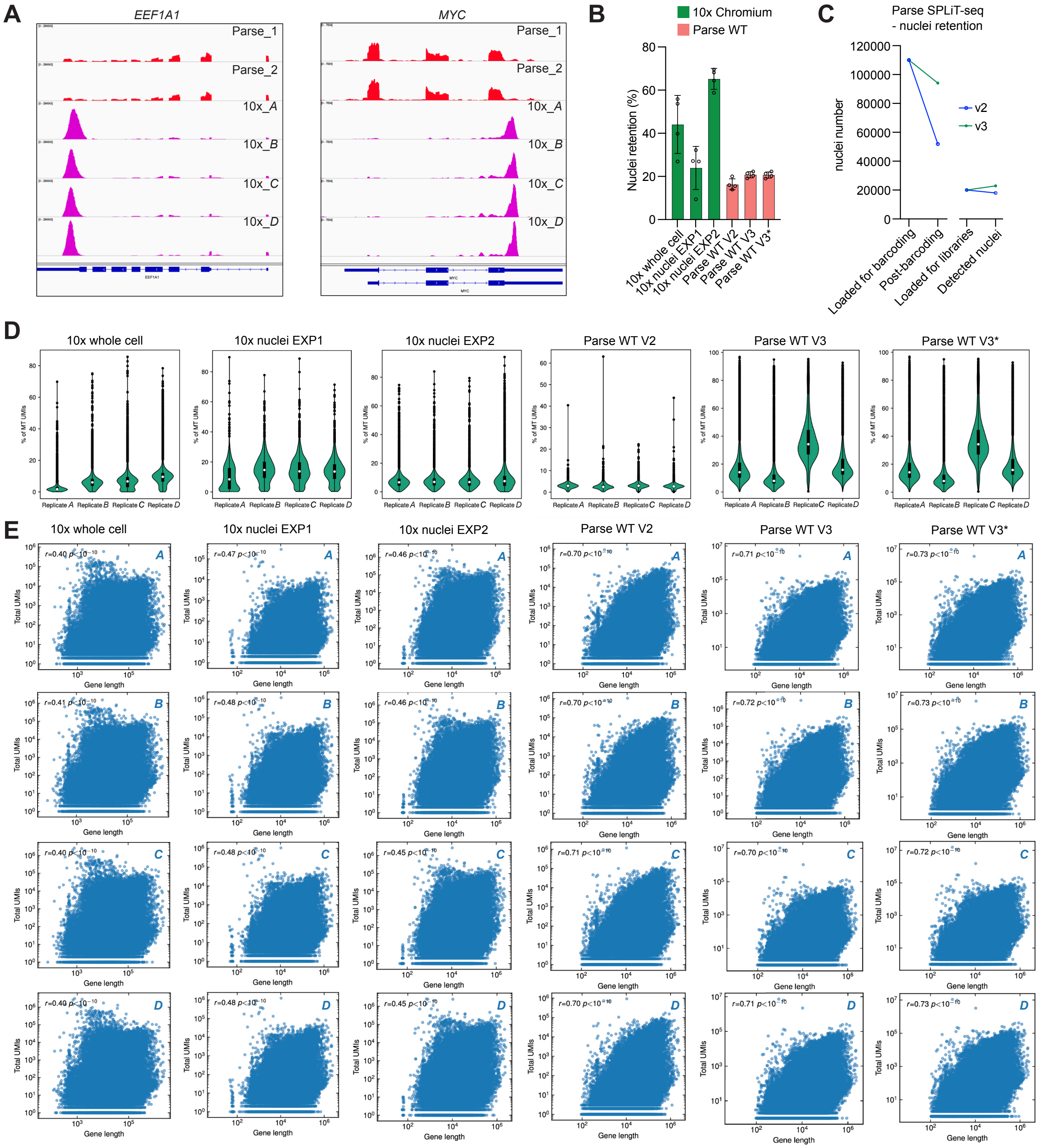


**Supplementary Figure S2. Sequencing library efficiencies related to Figure 2. (A)** Representative tracks of bulk single-nucleus RNA-seq shown for each replicate/library. The top panel shows the tracks for the *MYC* gene, and the bottom panel shows the *EEF1A1* gene as examples. Parse SPLiT-seq reads (library Parse_1 and library Parse_2) show even coverage of exonic sequences, while Chromium 10x (libraries *A*, *B*, *C*, and *D*) shows, as expected, 3' end-enriched reads. These results align with the use of oligo(dT) primers for cDNA synthesis in the 10x Chromium platform and both oligo dT and random hexamers in Parse technology. **(B)** Nuclei retention is shown as a percentage of recovered nuclei per experiment. Circles represent individual replicate values, bars show mean values, and error bars indicate SD. **(C)** Number of cells retained at different steps of the Parse SPLiT-seq protocol. *Loaded for barcoding* — number of nuclei in the starting material. Briefly, 109,000 are loaded for a yield of 20,000 cells. *Post-barcoding* - number of cells detected after three rounds of barcoding. *Loaded for libraries* - number of cells used to generate two sequencing libraries, *i.e.,* 10,000 cells in each library. *Detected nuclei* - final yield of nuclei from the experiment (see also Table 1). **(D)** Violin plots showing mitochondrial gene fractions as a proportion of total UMIs detected across separate replicates for all experiments. **(E)** Scatterplots of gene length versus the number of UMIs for all replicates and experiments, with '*r*' indicating the correlation between gene length and gene expression. Letters in italics *A*, *B*, *C,* and *D* designate individual replicates. See also main Figure 2.

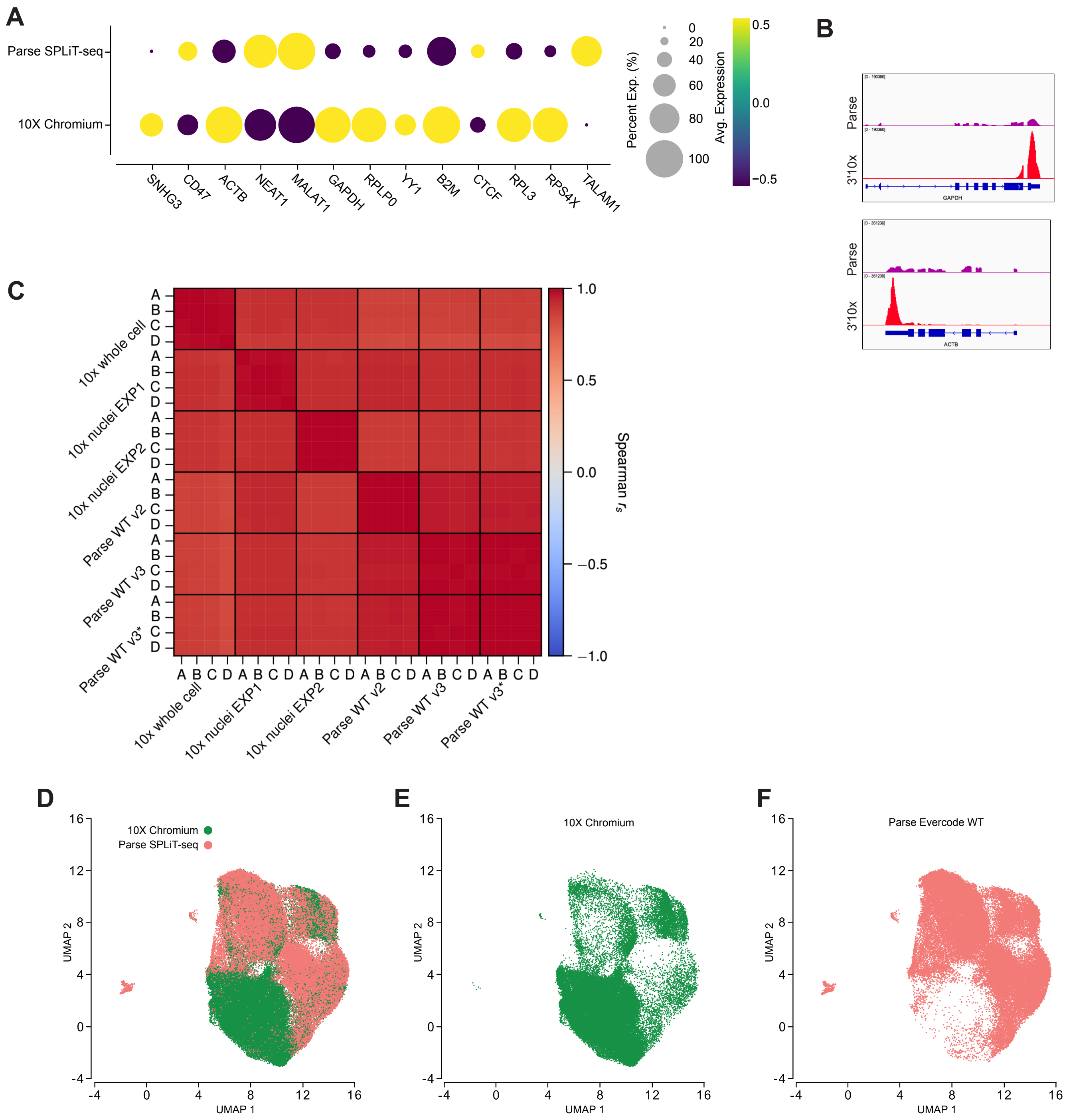


**Supplementary Figure S3. Platform-specific differences in biological parameters. (A)** Dot plot depicting the expression of selected genes (*SNHG3, CD47, ACTB, NEAT1, MALAT1, GAPDH, RPLP0, YY1, B2M, CTCF, RPL3, RPS4X, TALAM1*) comparing the combined experiments of Parse SPLIT-seq and 10X Chromium platforms. Circle size represents the percentage of cells expressing a gene (Percent Exp. (%)), and color intensity indicates average expression levels (Avg. Expression). **(B)** Representative genomic coverage tracks comparing the gene coverage of two housekeeping genes, *GAPDH* and *ACTB*, in Parse and 3' 10x platforms. **(C)** Pairwise Spearman correlation matrix ($r_s$) of gene expression across experiments (10x whole cell, 10x nuclei EXP1, 10x nuclei EXP2, Parse WT v2, Parse WT v3, and Parse WT v3*) and technical replicates (*A, B, C*, and *D*). **(D–F)** UMAP projection of single nuclei colored by platform. Overlapping cells from both platforms are shown in (D); cells detected by 10x Chromium are shown as green dots in panel (E); Parse Evercode WT is shown in (F).

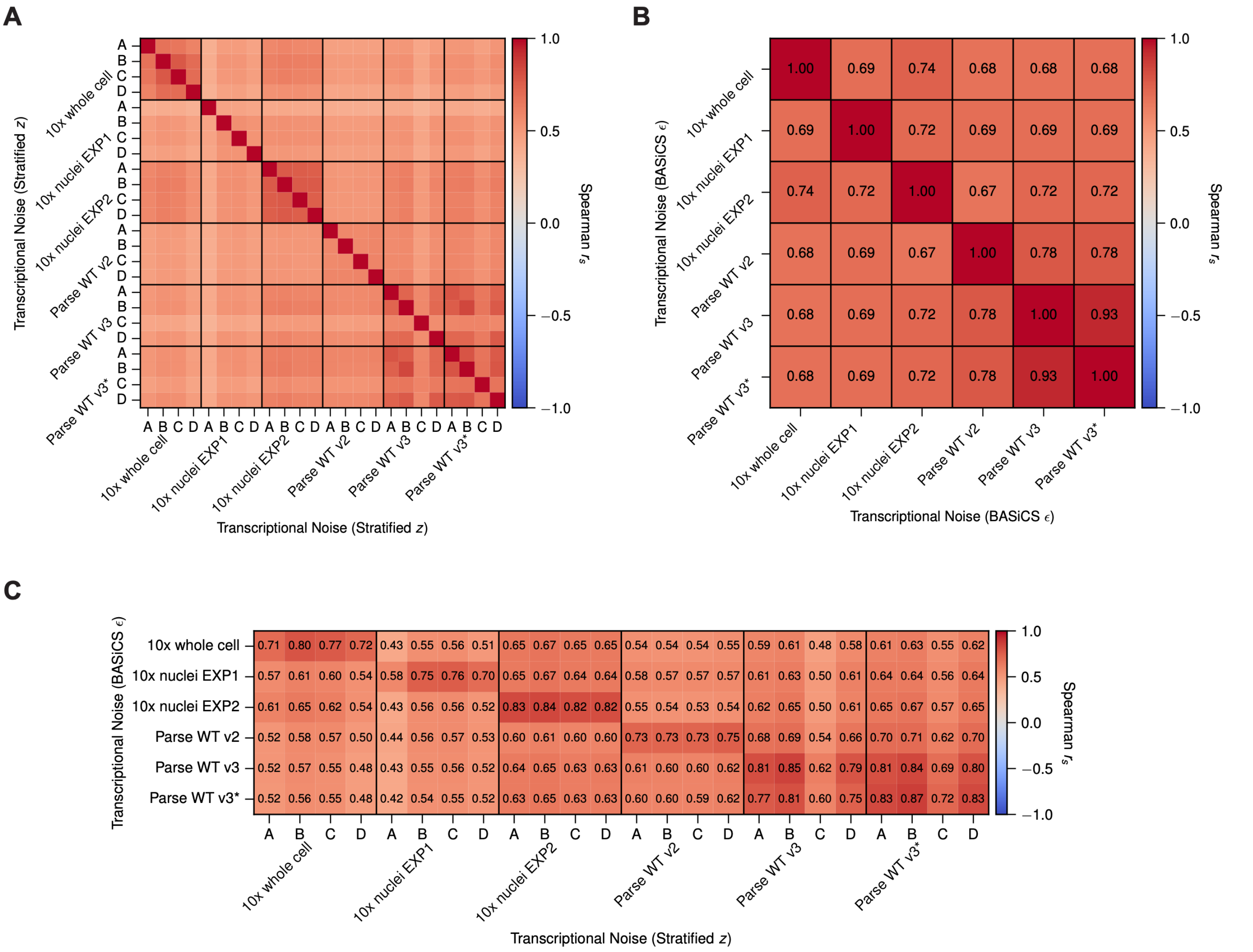


**Supplementary Figure S4. Reproducibility of transcriptional noise estimates within and between experiments. (A)** Spearman rank correlation of the stratified *z*-score between all pairs of individual replicates. Replicates (denoted by letters) are ordered by experiment, which are separated by black lines. Correlations were computed over the common set of genes detected in all six experiments. **(B)** Spearman rank correlation of the BASiCS residual overdispersion  between experiments. Since BASiCS estimates a single set of parameters per experiment, with replicates included as batches, no within-experiment comparison is available, and diagonal entries are self-correlations. **(C)** Spearman rank correlation between the BASiCS  of each experiment (rows) and the stratified *z*-score of each individual replicate (columns).

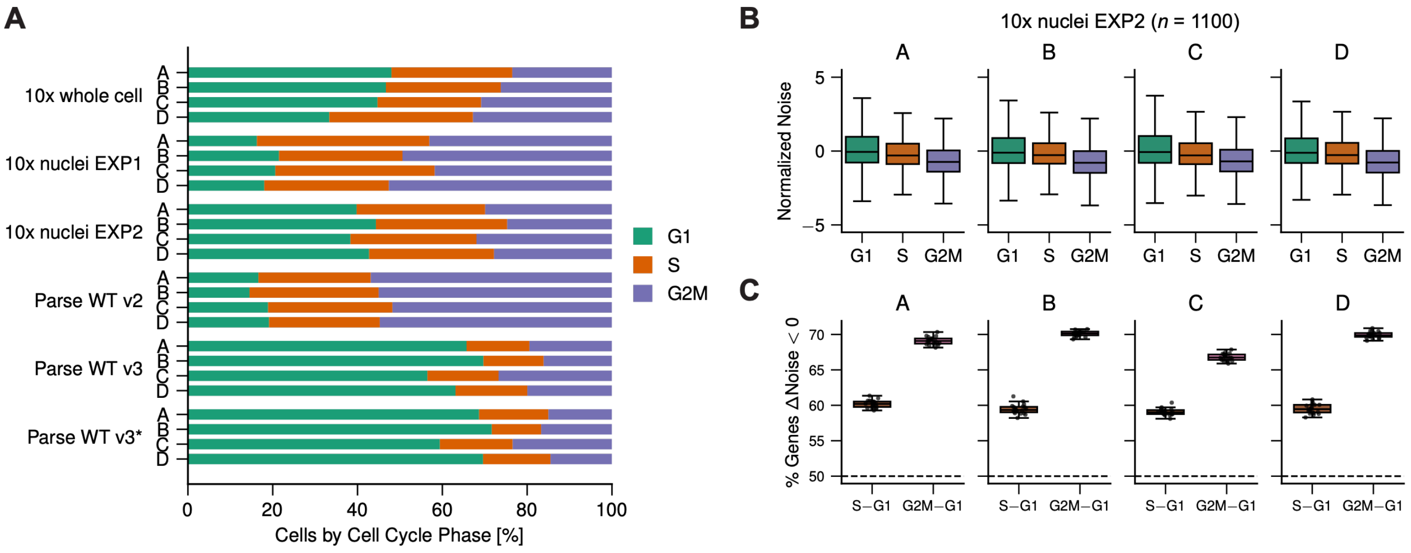


**Supplementary Figure S5. Cell-cycle phase composition and noise comparisons for individual replicates. (A)** Relative proportion of cells in each cell-cycle phase, shown for each replicate separately. Letters denote replicates within an experiment. The corresponding replicate-averaged proportions are shown in **Figure 4E**. **(B)** Distributions of transcriptional noise by cell-cycle phase for each replicate of 10x nuclei EXP2, with cells subsampled to  per phase. Boxes show the interquartile range and median, with whiskers extending to 1.5× the interquartile range and outliers not shown. **(C)** Proportion of genes with reduced noise in S or G2M relative to G1, for each replicate, over 20 independent subsamples of  cells per phase. Each point is one subsample, and the dashed line marks the 50% expected under no difference. Genes with identical estimates in both phases were counted in neither direction.

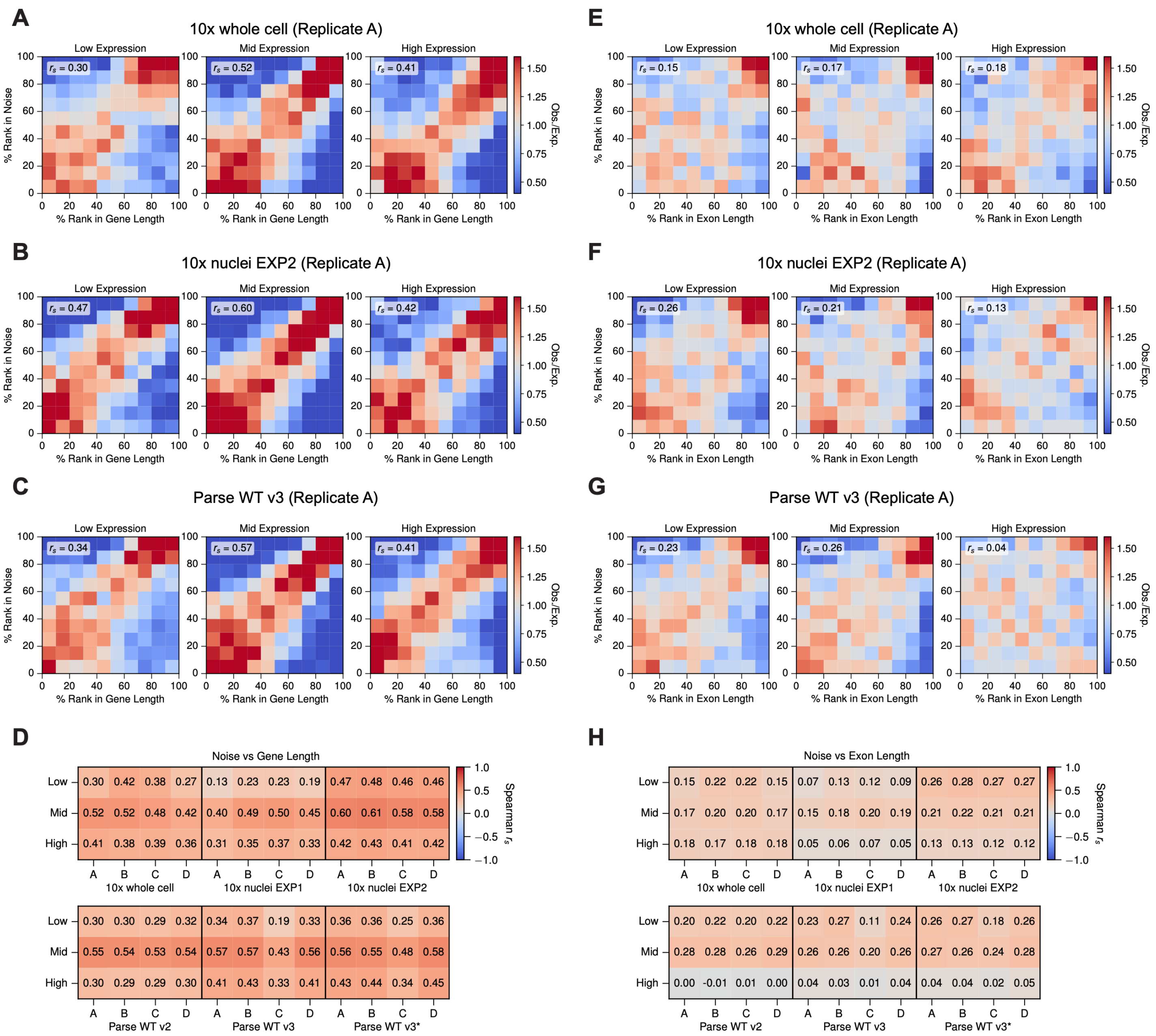


**Supplementary Figure S6. Association between transcriptional noise and gene length or exon length, based on the stratified *z*-score. (A–C)** Joint distribution of transcriptional noise and gene length for 10x whole cell (A), 10x nuclei EXP2 (B), and Parse WT v3 (C), shown separately for genes in the low, mid, and high level of mean expression. Genes were divided into deciles along each axis, and bin color gives the observed number of genes relative to that expected under independence. Spearman rank correlation between noise and gene length is shown for each expression stratum. Data are from Replicate A of each experiment, and panel B corresponds to **Figure 5A**. **(D)** Spearman rank correlation between noise and gene length for each replicate and expression stratum, grouped by experiment. Letters denote replicates within an experiment. The replicate-averaged values are shown in **Figure 5B**. **(E–H)** As in (A–D), using total exon length in place of genomic span. Panel F corresponds to **Figure 5C**, and the replicate-averaged values in (H) to **Figure 5D**. Correlations were computed over the common set of genes detected in all six experiments.

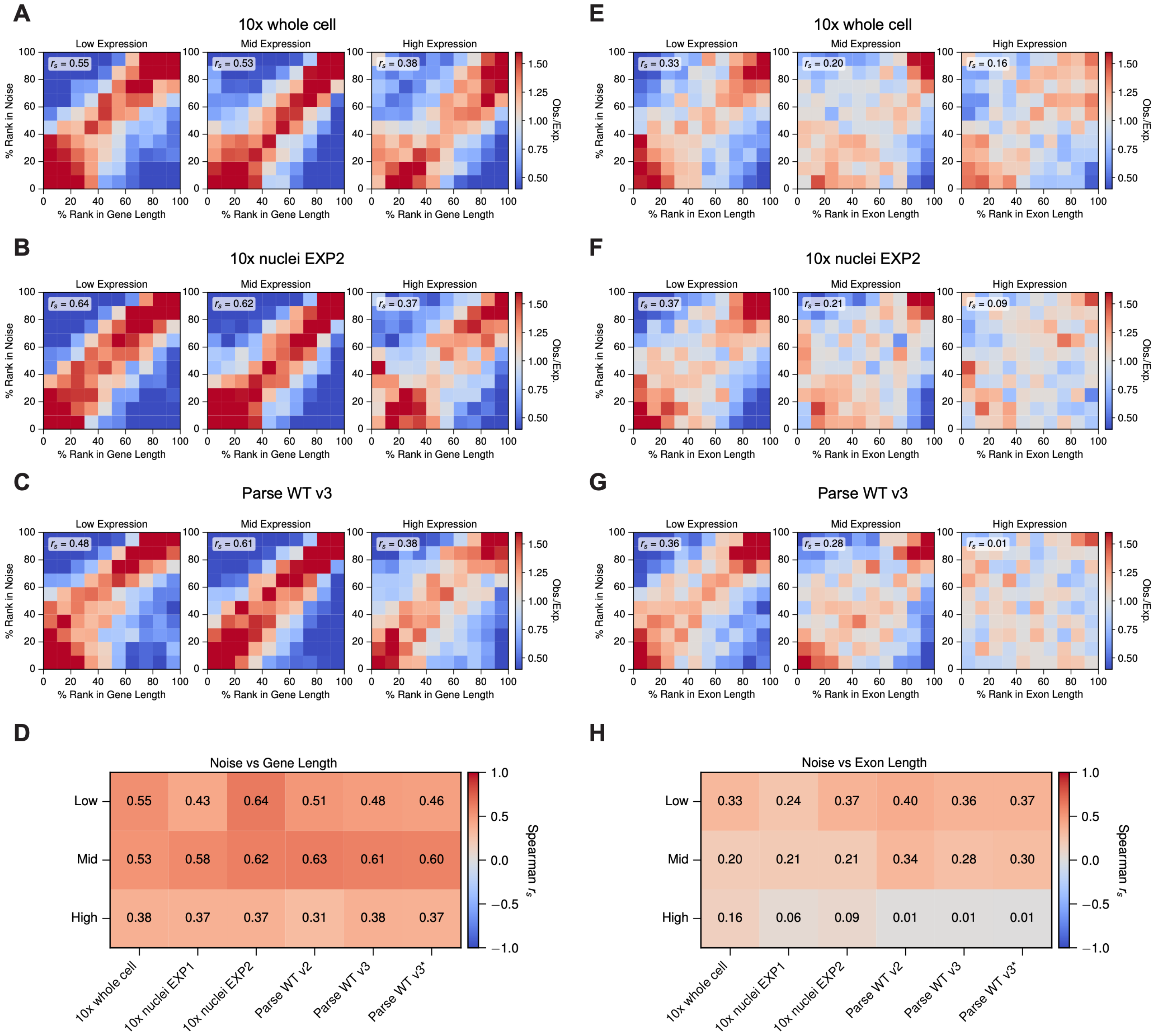


**Supplementary Figure S7. Association between transcriptional noise and gene length or exon length, based on the BASiCS residual overdispersion . (A–C)** Joint distribution of transcriptional noise and gene length for 10x whole cell (A), 10x nuclei EXP2 (B), and Parse WT v3 (C), shown separately for genes in the low, mid, and high level of mean expression. Genes were divided into deciles along each axis, and bin colour gives the observed number of genes relative to that expected under independence. Spearman rank correlation  between noise and gene length is shown for each expression stratum. **(D)** Spearman rank correlation between noise and gene length for each experiment and expression stratum. Since BASiCS estimates a single set of parameters per experiment, with replicates included as batches, one value was obtained per experiment rather than per replicate. **(E–H)** As in (A–D), using total exon length in place of genomic span. Correlations were computed over the common set of genes detected in all six experiments.

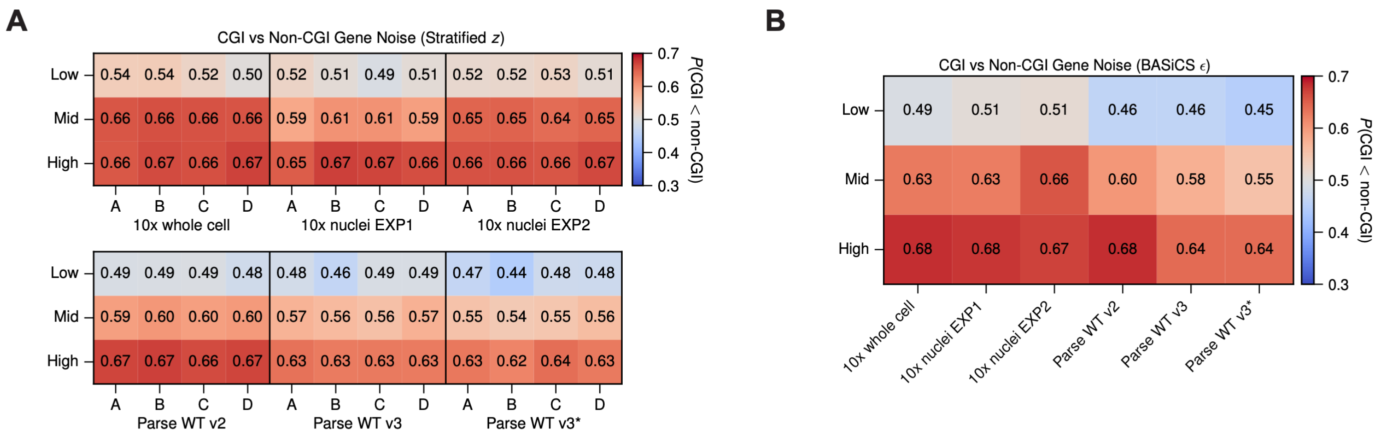


**Supplementary Figure S8. Reduced transcriptional noise in genes with CpG island promoters, across experiments and replicates. (A)** Probability that a randomly chosen CGI-promoter gene (CGI gene) has lower noise than a randomly chosen non-CGI gene, based on the stratified *z*-score, shown for each replicate and expression stratum. Letters denote replicates within an experiment. Values above 0.5 indicate reduced noise in CGI genes. The replicate-averaged values are shown in **Figure 5F**. **(B)** As in (A), based on the BASiCS residual overdispersion . Since BASiCS estimates a single set of parameters per experiment, with replicates included as batches, one value was obtained per experiment rather than per replicate. Genes were classified as CGI genes if their transcription start site fell within 1 kbp of an annotated CpG island, and comparisons were made over the common set of genes detected in all six experiments.
